# Physiological and metabolic insights into the first cultured anaerobic representative of deep-sea *Planctomycetes* bacteria

**DOI:** 10.1101/2023.07.18.549452

**Authors:** Rikuan Zheng, Chong Wang, Rui Liu, Ruining Cai, Chaomin Sun

## Abstract

*Planctomycetes* bacteria are ubiquitously distributed across various biospheres and play key roles in global element cycles. However, few deep-sea *Planctomycetes* members have been cultivated, limiting our understanding of *Planctomycetes* in the deep biosphere. Here, we have successfully cultured a novel strain of *Planctomycetes* (strain ZRK32) from a cold seep sediment, by using an enriched medium supplemented with rifampicin and different nitrogen sources. Our genomic, physiological and phylogenetic analyses indicate that strain ZRK32 is a novel species, which we propose be named: *Poriferisphaera heterotrophicis*. We show that strain ZRK32 replicates using a budding mode of division. Based on the combined results from growth assays and transcriptomic analyses, we found that rich nutrients, or supplementation with NO_3_^-^ or NH_4_^+^ promoted the growth of strain ZRK32 by facilitating energy production through the tricarboxylic acid (TCA) cycle and the Embden–Meyerhof–Parnas (EMP) glycolysis pathway. Moreover, supplementation with NO_3_^-^ or NH_4_^+^ induced strain ZRK32 to release a bacteriophage in a chronic manner, without host cell lysis. This bacteriophage then enabled strain ZRK32, and another marine bacterium that we studied, to metabolize nitrogen through the function of auxiliary metabolic genes (AMGs). Overall, these findings expand our understanding of deep-sea *Planctomycetes* bacteria, while highlighting their ability to metabolize nitrogen when reprogrammed by chronic viruses.

## Introduction

*Planctomycetes* bacteria are ubiquitous in many environments, including lakes (Pollet et al., 2011), wetlands (Dedysh and Ivanova, 2019), soil (Buckley et al., 2006), freshwater (Brümmer et al., 2004), oceanic waters, and abyssal sediments (Woebken et al., 2007; Goffredi and Orphan, 2010), where they are critical for carbon and nitrogen cycling; although, the specific pathways used are unknown (Wiegand et al., 2018). Although *Planctomycetes* bacteria are highly abundant in nature, relatively few have been cultivated; therefore, the unexplored groups lack cultured and characterized representatives (Fuerst and Sagulenko, 2011). Currently, all taxonomically described *Planctomycetes* with validly published names can be divided into two recognized classes: *Planctomycetia* and *Phycisphaerae* (Fukunaga et al., 2009; Wiegand et al., 2018). *Candidatus* Brocadiae is considered a third class within the phylum *Planctomycetes*, although pure cultures are not yet available for its members and its class-level status remains to be defined (Kartal et al., 2012; Kartal et al., 2013). The majority of the species isolated from the phylum *Planctomycetes* are from the class, *Planctomycetia* — a class that consists of four orders: *Planctomycetales*, *Pirellulales*, *Gemmatales*, and *Isosphaerales* (Dedysh et al., 2020). In contrast, only few members from the *Phycisphaerae* class have been cultured; the class *Phycisphaerae* contains three orders: *Phycisphaerales* (Fukunaga et al., 2009), *Tepidisphaerales* (Kovaleva et al., 2015), and *Sedimentisphaerales* (Spring et al., 2018).

*Planctomycetes* bacteria have fascinating physiological characteristics (Wiegand et al., 2018). For decades, *Planctomycetes* bacteria have blurred the lines between prokaryotes and eukaryotes. *Planctomycetes* bacteria possess several uncommon traits when compared with typical bacteria: they have a compartmentalized cell plan, an enlarged periplasm, a tightly folded nucleus-like structure, an endocytosis-like method of uptake, and a FtsZ-free method of cell division (Fuerst and Webb, 1991; Lindsay et al., 1997; Lonhienne et al., 2010; Wiegand et al., 2018). The unique cellular structures in *Planctomycetes* bacteria have stretched and challenged our understanding of the concept of a ‘prokaryote’ (Fuerst and Sagulenko, 2011).

*Planctomycetes* bacteria also exhibit a diverse range of respiration and cell division methods. Members of the class *Planctomycetia* are mostly aerobic and divide by budding, while *Phycisphaerae* members are mostly anaerobic and divide by binary fission (Rivas-Marín et al., 2016; Wiegand et al., 2018; Pradel et al., 2020; Wiegand et al., 2020b). However, a recent report showed that *Poriferisphaera corsica* KS4 — a novel strain from class *Phycisphaerae —* was aerobic and that it might divide by budding (Kallscheuer et al., 2020b); this is different from previous reports and suggests that there are many physiological and cellular *Planctomycetes* bacteria characteristics yet to be discovered.

The deep sea is where life may have originated from and where stepwise evolution occurred (Orgel, 1998); it is also where a large number of uncultured microorganisms live (Zheng et al., 2021b). Among these microbes, *Planctomycetes* might dominate deep-sea sediments (Wiegand et al., 2018). For example, in the Gulf of Mexico, the phylum *Planctomycetes* accounts for 28% of all bacteria, where they seem to be involved in the nitrogen cycle, and the breakdown of organic detrital matter, which is delivered to the sediment as marine snow (Vigneron et al., 2017). Unfortunately, only few *Planctomycetes* bacteria from deep-sea environments have been cultured (Wang et al., 2020; Wiegand et al., 2020b), limiting our understanding of their characteristics (e.g., material metabolism, element cycling, and ecological role) (Kulichevskaya et al., 2020).

Here, we successfully cultured a novel member of *Planctomycetes* (strain ZRK32) from a deep-sea subsurface sediment. We found that strain ZRK32 used a budding mode of division. We also found that supplementation with rich nutrients, and either NO_3_^-^ or NH_4_^+^, promoted strain ZRK32 growth. Moreover, the presence of NO_3_^-^ or NH_4_^+^ induced strain ZRK32 to release a bacteriophage in a chronic manner— a process that does not kill the host cell (i.e., the host bacteria continues to grow despite phage reproduction) (Liu et al., 2022). This bacteriophage reprogrammed strain ZRK32, and another marine bacterium we studied, to metabolize nitrogen through the action of auxiliary metabolic genes (AMGs).

## Results

### Isolation, morphology, and phylogenetic analysis of a novel strain of *Planctomycetes* isolated from a deep-sea cold seep

We enriched deep-sea *Planctomycetes* bacteria using a basal medium supplemented with rifampicin and inorganic nitrogen sources (NaNO_3_ and NH_4_Cl). These enriched samples were plated on to agar slants in Hungate tubes, and then colonies with distinct morphologies were selected and cultivated (Figure 1A). Some colonies were noted to be from the phylum *Planctomycetes*, based on their 16S rRNA sequences. Among these, strain ZRK32 was selected for further study as it grew faster than other strains. Following negative staining and TEM observation, we observed that strain ZRK32 cells were spherical (with an average diameter of 0.4-1.0 µm), and had a single polar flagellum (Figure 1B). Moreover, we found that the mother and daughter cells of strain ZRK32 had distinct sizes at the stage of cell division (Figure 1C and Figure S1), indicating that strain ZRK32 was dividing asymmetrically (i.e., through budding). Ultrathin whole-cell sections showed that strain ZRK32 possessed a condensed and intact nucleoid-like structure, and a complex extended membrane structure (Figure 1D).

**Figure 1.**
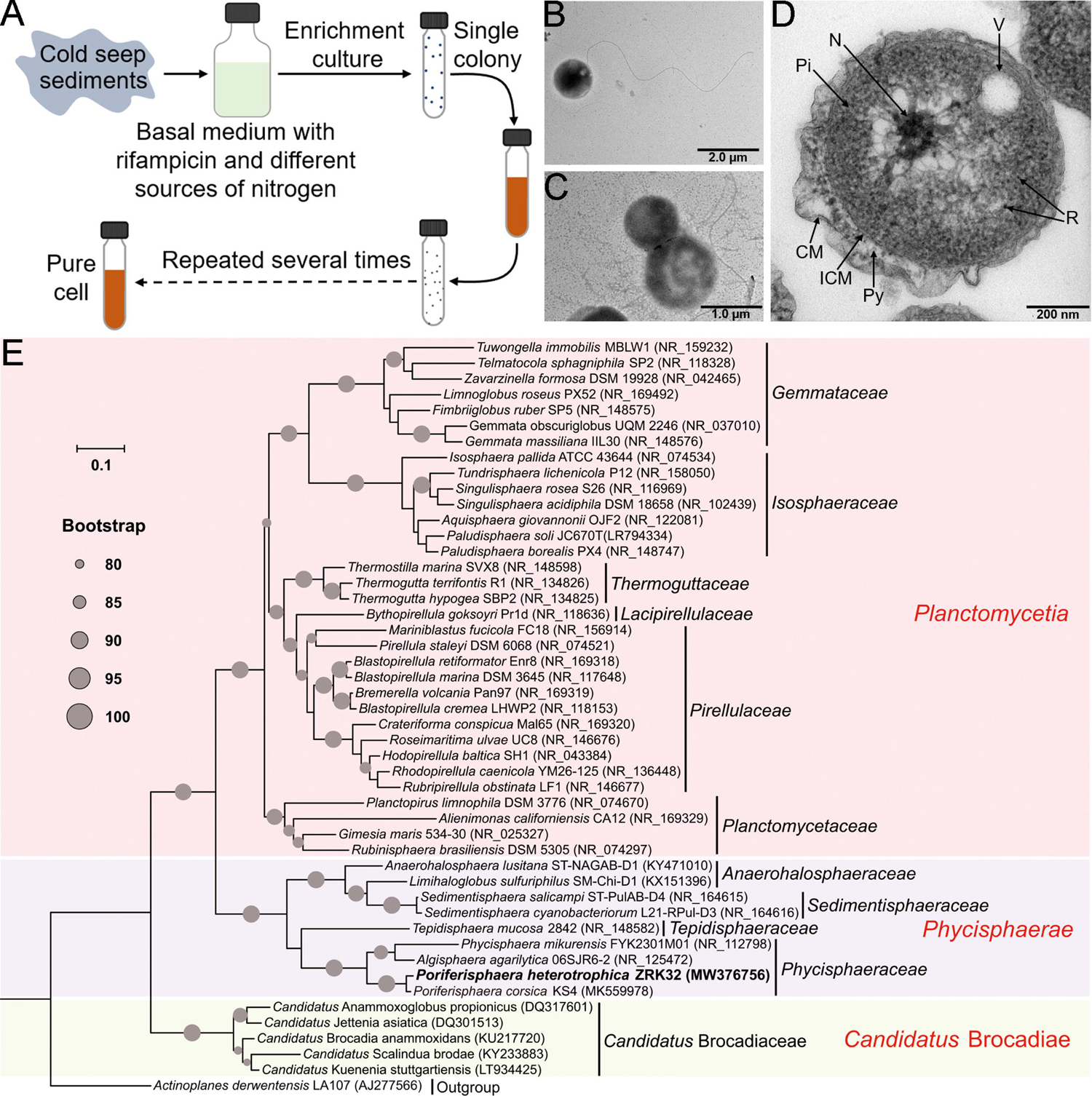
Isolation, morphology, and phylogenetic analysis of *P. heterotrophicis* ZRK32. (A) Diagram showing the strategy used to isolate the *Planctomycetes* bacteria. (B, C) TEM observation of strain ZRK32. (D) TEM observation of ultrathin sections of cells from strain ZRK32. Abbreviations: CM, outer membrane; Pi, cytoplasm; R, ribosome; N, nucleoid; ICM, cytoplasmic membrane; Py, periplasm; V, vesicle-like organelles. (E) Phylogenetic analysis of strain ZRK32. Phylogenetic placement of strain ZRK32 within the phylum *Planctomycetes*, based on almost complete 16S rRNA gene sequences. The NCBI accession number for each 16S rRNA gene is indicated after each corresponding strain’s name. The tree was inferred and reconstructed using the maximum likelihood criterion, with bootstrap values (%) > 80; these are indicated at the base of each node with a gray dot (expressed as a percentage from 1,000 replications). The 16S rRNA gene sequence of *Actinoplanes derwentensis* LA107^T^ was used as the outgroup. Bar, 0.1 substitutions per nucleotide position.

Based on the 16S rRNA sequence of strain ZRK32, a sequence similarity calculation using the NCBI server indicated that the closest relatives of strain ZRK32 were *Poriferisphaera corsica* KS4^T^ (98.06%), *Algisphaera agarilytica* 06SJR6-2^T^ (88.04%), *Phycisphaera mikurensis* NBRC 102666^T^ (85.28%), and *Tepidisphaera mucosa* 2842^T^ (82.94%). Recently, the taxonomic threshold for species based on 16S rRNA gene sequence identity value was 98.65% (Kim et al., 2014). Based on these criteria, we proposed that strain ZRK32 might be a novel representative of the genus *Poriferisphaera*. In addition, to clarify the phylogenetic position of strain ZRK32, the genome relatedness values were calculated by the average nucleotide identity (ANI), the tetranucleotide signatures (Tetra), and *in silico* DNA-DNA similarity (*is*DDH), against the genomes of strains ZRK32 and KS4. The ANIb, ANIm, Tetra, and *is*DDH values were 72.89%, 85.34%, 0.97385, and 20.90%, respectively (Table S1). These results together demonstrated the strain ZRK32 genome to be obviously below established ‘cut-off’ values (ANIb: 95%, ANIm: 95%, Tetra: 0.99, *is*DDH: 70%) for defining bacterial species, suggesting strain ZRK32 represents a novel strain within the genus *Poriferisphaera*.

To further confirm the taxonomy of strain ZRK32, we performed phylogenetic analyses. The maximum likelihood tree of 16S rRNA indicated that strain ZRK32 was from the genus *Poriferisphaera* and that it formed an independent phyletic line with strain *Poriferisphaera corsica* KS4^T^ (Figure 1E). The genome tree also suggested that this novel clade was a sister-strain of strain KS4^T^, which belongs to the genus *Poriferisphaera* (Figure S2). Based on its genomic (Figure S3 and Table S1), physiological (Figure S4) and phylogenetic characteristics, strain ZRK32 was distinguishable from strain *Poriferisphaera corsica* KS4^T^, which is currently the only species of the genus *Poriferisphaera* with a validly published name. We therefore propose that strain ZRK32 represents a novel species in the genus *Poriferisphaera*, for which the name *Poriferisphaera heterotrophicis* sp. nov. is proposed.

### Rich nutrients promote *P. heterotrophicis* ZRK32 growth

The growth rate of strain ZRK32 increased when it was cultured in a rich medium (containing ten times more yeast extract than basal medium) (Figure 2A). To gain further insight into its metabolic characteristics, we performed transcriptomic analyses of strain ZRK32 grown in the rich medium and strain ZRK32 grown in the basal medium. The results showed that the expression of many genes involved in the TCA cycle (Figure 2B, C) and the EMP glycolysis pathway (Figure S5A, B) (which both contribute to energy production) were up-regulated. In addition, the expression of genes encoding NADH-ubiquinone oxidoreductase and flagellum assembly-related proteins were also up-regulated (Figure 2D, E). To verify the transcriptomic data, we performed qRT-PCR assays, which showed the same gene expression variation, consistent with the transcriptomic results (Figure S6). Based on the combined results of the growth assay and transcriptomic analyses, we concluded that strain ZRK32 growth is better when cultured using nutrient-rich medium.

**Figure 2.**
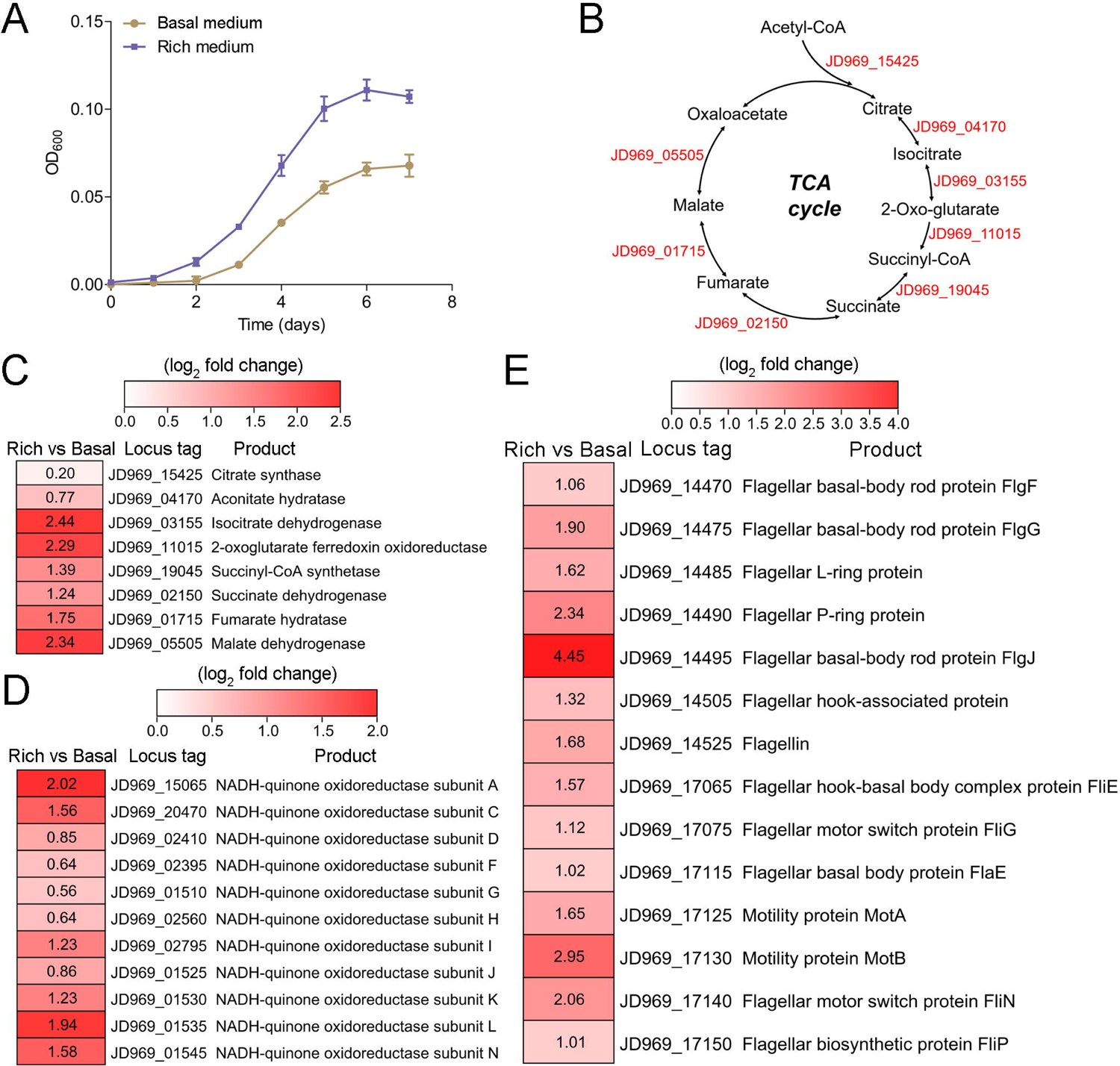
Growth assay and transcriptomic analysis of *P. heterotrophicis* ZRK32 strains cultivated in basal medium and rich medium. (A) Growth curves of ZRK32 strains cultivated in basal medium and rich medium. (B) Diagram of the TCA cycle. The gene numbers shown in this schemeatic are the same as those shown in panel C. Transcriptomics-based heat map showing the relative expression levels of genes associated with the TCA cycle (C), NADH-quinone oxidoreductase (D), and flagellar assembly (E) of strain ZRK32 cultivated in rich medium (Rich) compared with strain cutivated in basal medium (Basal). The numbers in panels C, D, and E represent the fold change of gene expression (by using the log_2_ value).

### *P. heterotrophicis* ZRK32 replicates using a budding mode of division

After reviewing more than 600 TEM photos, we confirmed that strain ZRK32 divided by budding —a method also reported in other *Planctomycetes* bacteria (Figure 1C and Figure S1) (Wiegand et al., 2020b). Remarkably, during the early stages of budding in strain ZRK32, the extracellular membrane extended and formed a bulge, which grew until it was a similar size to the mother cell (Figure 3A, B, panels 1 to 4). The genetic materials within the nucleoid then duplicated and divided equally between the mother and daughter cells, along with other cytoplasmic contents (Figure 3A, B, panels 5 to 8). The daughter cell then separated from the mother cell, completing cell division.

**Figure 3.**
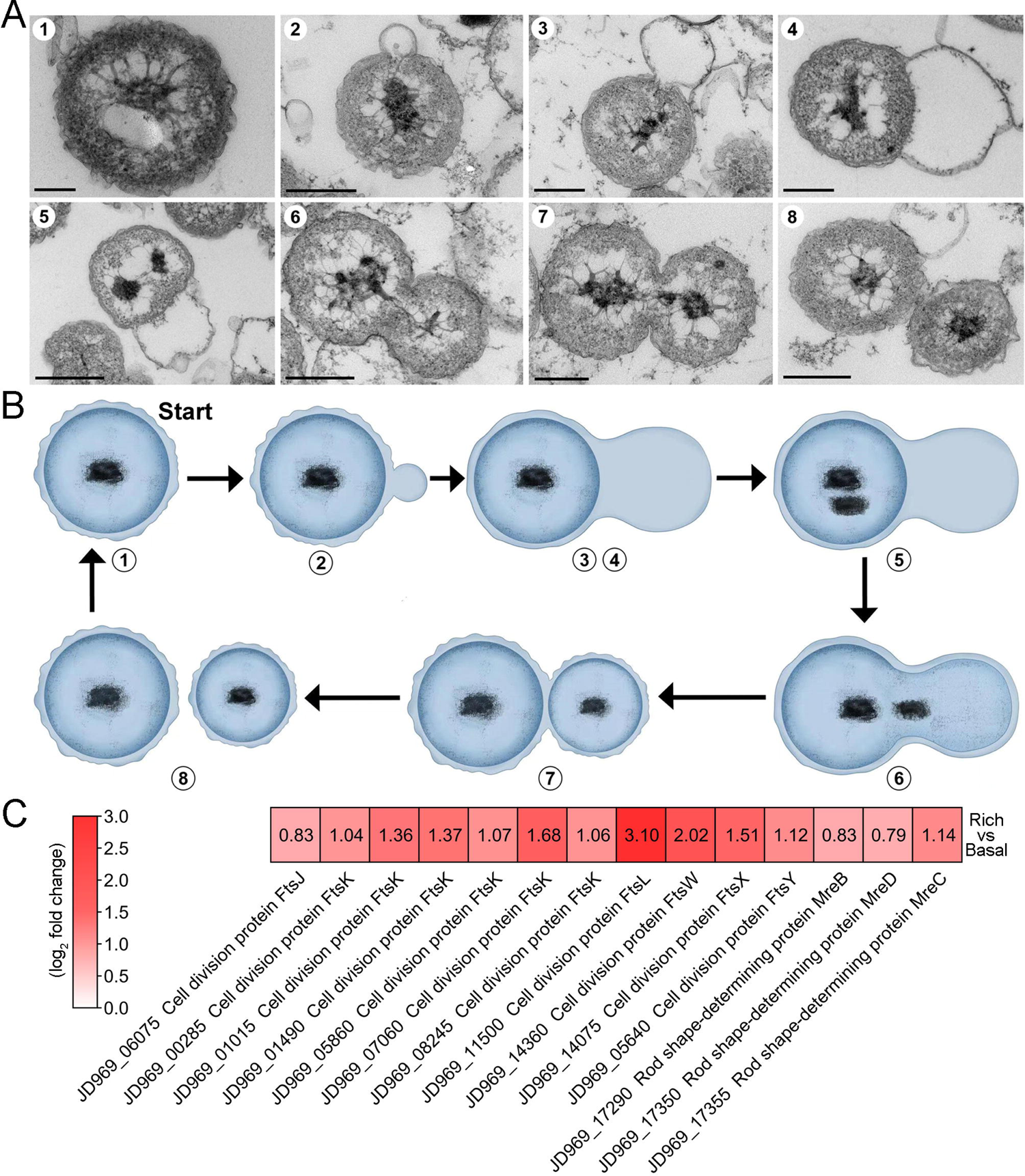
The mode of cell division utilized by *P. heterotrophicis* ZRK32. (A) Ultrathin TEM sections showing the process of polar budding division (panels 1-8) in strain ZRK32. Images representing the different phases of cell division are shown. Scale bars are 200 nm in panels A and B. (B) The proposed model of cell division of strain ZRK32 based on the TEM observation shown in panel B. The numbers in panels A and B correspond to the same phase of division. (C) Transcriptomics-based heat map showing the differentially expressed genes that encode different key proteins associated with cell division in strain ZRK32. The numbers in panel A represent the fold change of gene expression (by using the log_2_ value).

Next, we investigated whether genes associated with budding method of cell division were present in the genome of strain ZRK32, and whether they were functional during bacterial growth. We performed genomic and transcriptomic analyses of strain ZRK32. We did not detect the cell division protein, FtsZ, in strain ZRK32, but we did identify other Fts-related proteins (e.g., FtsJ, FtsK, FtsL, FtsW, FtsX and FtsY) and rod shape-determining proteins (MreB, MreC and MreD); the expressions of genes encoding these Fts-related proteins were upregulated in the ZRK32 strains cultured in the rich medium compared with strains cultured in basal medium (Figure 3C).

### Effects of NO_3_^-^, NH_4_^+^, and NO_2_^-^ on *P. heterotrophicis* ZRK32 growth

As *Planctomycetes* bacteria are involved in nitrogen cycling, we tested the effects of different nitrogen-containing substances (including NO_3_^-^, NH_4_^+^ and NO_2_^-^) on strain ZRK32 growth. These assays showed that adding NO_3_^-^ or NH_4_^+^ to the culture medium increased strain ZRK32 growth, while adding NO_2_^-^ inhibited growth (Figure 4A). The concentration of NO_3_^-^ decreased from ∼21 mM to ∼6 mM, and then to ∼4 mM after strain ZRK32 had been incubating for four days and six days, respectively. The concentration of NH_4_^+^ increased from ∼0 mM to ∼11 mM, and then to ∼7 mM after strain ZRK32 had been incubating for four days and six days, respectively. These results strongly suggest that strain ZRK32 can effectively convert NO_3_^-^ to NH_4_^+^ (Figure 4B). In addition, when strain ZRK32 was incubated in the rich medium supplemented with NH_4_ ^+^ for four days and six days, the concentration of NH_4_ ^+^ decreased from ∼19 mM to ∼11 mM, and then to ∼4 mM, with no change in the concentrations of NO_3_^-^ and NO_2_^-^ (Figure 4C). To investigate nitrogen metabolism in strain ZRK32, we analyzed the strain ZRK32 genome and found that it contained a complete nitrate reduction pathway and key genes responsible for the conversion of ammonia to glutamate (Figure 4D), which explains the results in figures 4B and C. Subsequently, we performed transcriptome sequencing analysis and found that the genes encoding nitrate reductase (NapA and NapB), nitrite reductase (NirB), glutamine synthetase, and glutamate synthase were all simultaneously up-regulated in the presence of NO_3_^-^. We also found that the genes encoding glutamine synthetase and glutamate synthase were up-regulated in the presence of NH_4_^+^ (Figure 4E). However, no differential expression was observed for the gene, *nirD* in the presence of NO_3_^-^ or NH_4_^+^, even though this gene encodes a nitrite reductase. Moreover, we observed that some genes involved in the TCA cycle (Figure 4F), the EMP glycolysis pathway (Figure S5C), and genes encoding NADH-ubiquinone oxidoreductase-related proteins (Figure 4G) were upregulated in the presence of NO_3_ ^-^ or NH_4_ ^+^, but down-regulated in the presence of NO_2_^-^. We also observed that the expressions of many genes associated with flagellum assembly were up-regulated when NO_3_ ^-^ or NH_4_ ^+^ were supplemented into the culture medium (Figure 4H). We observed similar trends in our qRT-PCR results (Figure S7), which validated our RNA-seq results.

**Figure 4.**
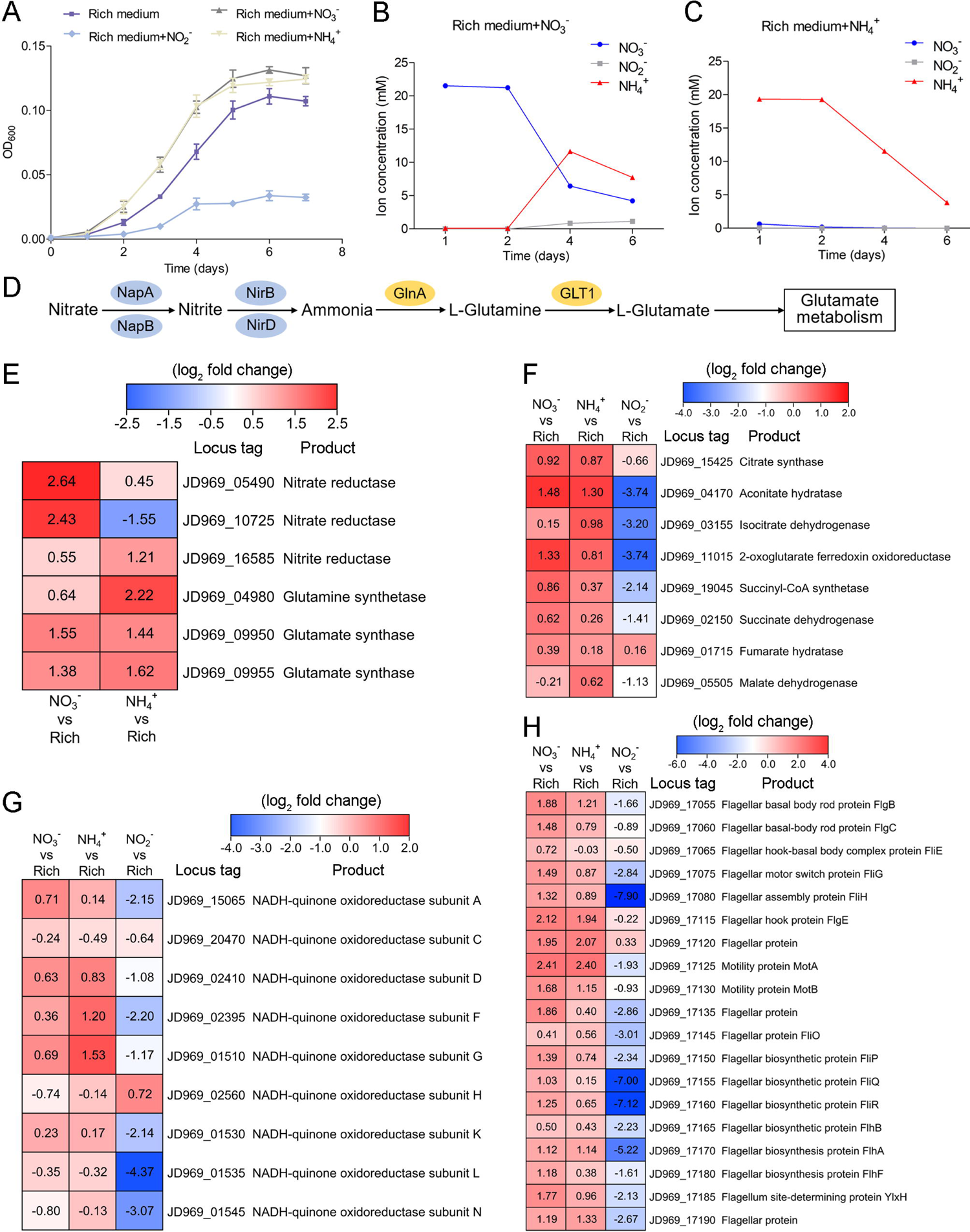
Nitrogen metabolism assays of *P. heterotrophicis* ZRK32. (A) Growth curves of ZRK32 strains cultivated in the rich medium alone and cultivated in rich medium supplemented with either 20 mM NO_3_^-^, 20 mM NH_4_^+^, or 20 mM NO ^-^. (B) The dynamics of concentrations of NO_3_^-^, NH_4_^+^, and NO_2_^-^ in strains of ZRK32 cultivated in the rich medium supplemented with 20 mM NO_3_^-^. (C) The dynamics of concentrations of NO_3_^-^, NH_4_^+^, and NO_2_^-^ in strains of ZRK32 cultivated in the rich medium supplemented with 20 mM NH ^+^. (D) The predicted nitrogen metabolism pathway of strain ZRK32. Abbreviations: NapA, periplasmic nitrate reductase; NapB, periplasmic nitrate reductase, electron transfer subunit; NirB, nitrite reductase (NADH) large subunit; NirD, nitrite reductase (NADH) small subunit; GlnA, glutamine synthetase; GLT1, glutamate synthase. Transcriptomics-based heat map showing the relative expression levels of genes associated with nitrogen metabolism (E), the TCA cycle (F), NADH-quinone oxidoreductase (G), and flagellar assembly (H) in strains of ZRK32 cultivated in the rich medium supplemented with different inorganic nitrogen sources (20 mM NO ^-^, 20 mM NH ^+^ or 20 mM NO ^-^) compared with strains cultivated in the rich medium alone. “Rich” indicates rich medium. “NO_3_^-^, NH_4_^+^, and NO ^-^” indicate rich medium supplemented with 20 mM NO ^-^, 20 mM NH ^+^, and 20 mM NO ^-^, respectively. The numbers in panels E, F, G, and H represent the fold change of gene expression (by using the log_2_ value).

### NO_3_^-^ and NH_4_^+^ induce the release of a chronic bacteriophage in *P. heterotrophicis* strain ZRK32

Bacteriophages are widely distributed across oceans and can regulate nitrogen metabolism in their host (Cassman et al., 2012; Monier et al., 2017; Gazitúa et al., 2021; Wang et al., 2022). We therefore investigated whether bacteriophages affected nitrogen metabolism in strain ZRK32. TEM observations showed that phage-like structures (hexagonal phages, ∼30 nm) were present in cell suspensions of the ZRK32 strain that had been cultured using nutrient-rich medium supplemented with either NO_3_^-^ or NH_4_^+^ (Figure 5A, panels II and III). In contrast, no phage-like structures were observed in the cell suspensions of ZRK32 strain that was cultured in the rich medium alone (without NO_3_^-^ or NH_4_^+^) (Figure 5A, panel I). This suggests that the presence of NO_3_^-^ or NH_4_^+^ stimulated the release of the bacteriophages from strain ZRK32. Most chronic bacteriophages do not negatively affect their host’s growth when cultivated in the laboratory (Alarcón-Schumacher et al., 2022). Consistently, the replication and release of the bacteriophages from strain ZRK32 did not kill the host cell, consistent well with the key feature of chronic bacteriophages (Howard-Varona et al., 2017). By comparing genomic sequences, we confirmed that the genome of the phage induced by NO_3_^-^ was the same as the phage induced by NH_4_^+^. When we compared this phage genome (Phage-ZRK32, 21.9 kb) (Figure 5B) with the host genome (strain ZRK32) (using Galaxy Version 2.6.0 (https://galaxy.pasteur.fr/) (Afgan et al., 2018) with the NCBI BLASTN method), we found that the Phage-ZRK32 genome was outside of the host chromosome; this indicates that this chronic bacteriophage is extrachromosomal, which is consistent with previous reports (Chevallereau et al., 2022). In addition to the genes encoding phage-associated proteins, the genome of Phage-ZRK32 also has many auxiliary metabolic genes (AMGs), which encode glutamine amidotransferase, amidoligase, glutathione synthase, and gamma-glutamylcyclotransferase (Figure 5B). Although, some genes (including genes encoding amidoligase, glutathione synthase and gamma-glutamylcyclotransferase) were absent from the strain ZRK32 genome.

**Figure 5.**
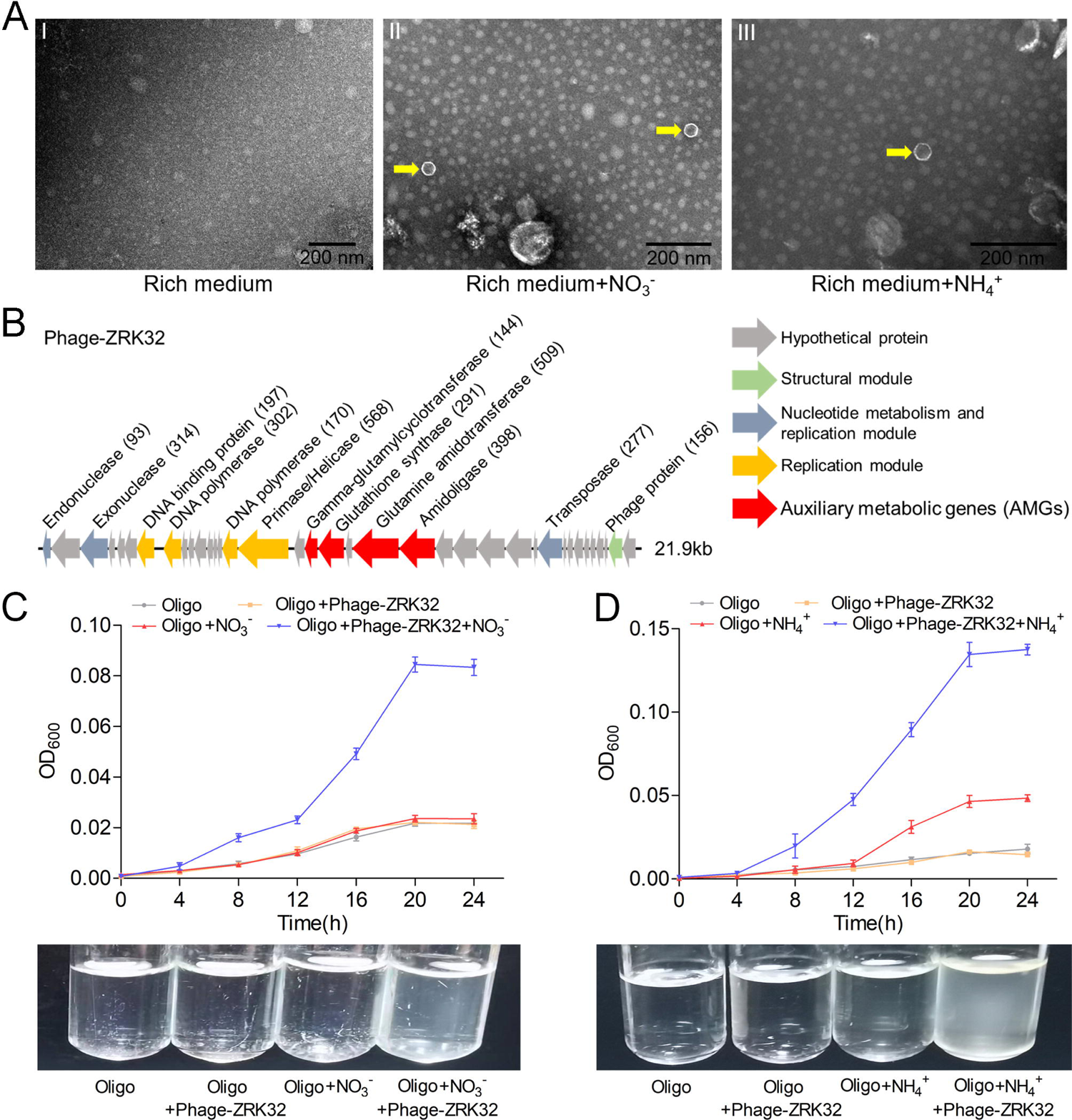
Observation and functional assay of the chronic bacteriophage induced by NO_3_^-^ or NH_4_^+^ from *P. heterotrophicis* ZRK32. (A) TEM observation of phages extracted from the cell suspensions of ZRK32 strains that cultured in either the rich medium alone, or rich medium supplemented with 20 mM of either NO_3_^-^ or NH_4_^+^. (A, Panel I) No phage-like particles were observed in the cell suspensions from the ZRK32 strain cultured in the rich medium. (A, Panels II and III) Hexagonal phages (indicated with yellow arrows) observed in the cell suspensions from the ZRK32 strains cultured in the rich medium supplemented with 20 mM of either NO_3_^-^ or NH_4_^+^. The scale bar is 200 nm. (B) A diagram showing the genomic composition of Phage-ZRK32. The arrows represent different ORFs and the direction of transcription. The main putative gene products of this phage are shown, and the numbers in brackets indicate the numbers of amino acids. Hypothetical proteins are indicated by gray arrows, structural modules are indicated by green arrows, nucleotide metabolism is indicated by blue-gray arrows, the replication module is indicated by gold arrows, and AMGs are indicated by red arrows. The size of the phage genome is shown beside the gene cluster. (C) Bacterial growth curve showing the growth rate of strains of *Pseudomonas stutzeri* 273 cultivated in either oligotrophic medium, oligotrophic medium supplemented with Phage-ZRK32, oligotrophic medium supplemented with 20 mM NO_3_^-^, or oligotrophic medium supplemented with 20 mM NO_3_^-^ and Phage-ZRK32. (D) Bacterial growth curve showing the growth rate of strains of *Pseudomonas stutzeri* 273 cultivated in either oligotrophic medium, oligotrophic medium supplemented with Phage-ZRK32, oligotrophic medium supplemented with 20 mM NH_4_^+^, or oligotrophic medium supplemented with 20 mM NH_4_^+^ and Phage-ZRK32. “Oligo” indicates oligotrophic medium.

To verify whether these AMGs were functional, we investigated whether Phage-ZRK32 was capable of reprogramming nitrogen metabolism and promoting growth in other marine bacteria. We selected the aerobic marine bacterium, *Pseudomonas stutzeri* 273 (Wu et al., 2016), and examined the effects of Phage-ZRK32 on its growth. This showed that Phage-ZRK32 promoted *Pseudomonas stutzeri* 273 growth by facilitating the metabolism and utilization of NO_3_^-^ and NH_4_^+^ (Figure 5C, D). In particular, adding Phage-ZRK32 and NH_4_^+^ into the oligotrophic medium resulted in an approximate 3–8-fold increase in growth compared with strains cultivated without NH_4_^+^ or Phage-ZRK32 supplementation (Figure 5D). The AMGs that encode amidoligase and gamma-glutamylcyclotransferase were absent from the *Pseudomonas stutzeri* 273 genome, even though it contains a complete nitrate reduction pathway and some genes responsible for converting ammonia to glutamate. Therefore, the Phage-ZRK32 AMGs might facilitate nitrogen metabolism and amino acid generation in *Pseudomonas stutzeri* 273 in a similar way to *P. heterotrophicis* ZRK32.

## Discussion

Until recently, most research on *Planctomycetes* has focused on strains in freshwater and shallow ocean environments (Bondoso et al., 2011; Kallscheuer et al., 2020b), with few studies on deep-sea strains; this is likely due to the logistical difficulties associated with sampling and cultivating these strains. The availability of cultured and characterized representatives for many phylogenetic clades within the phylum *Planctomycetes* are lacking (Dedysh et al., 2021). Therefore, more approaches and media types, like those used in our study (Figure 1A), should be developed to obtain *Planctomycetes* bacteria from different deep-sea environments. The vast majority of *Planctomycetes* members have been isolated using an oligotrophic medium supplemented with N-acetyl glucosamine (Kaboré et al., 2020; Kallscheuer et al., 2020a; Peeters et al., 2020; Salbreiter et al., 2020; Wiegand et al., 2020a); use of this medium previously resulted in the breakthrough isolation of 79 *Planctomycetes* strains (Wiegand et al., 2020b). However, we observed that strains of *P. heterotrophicis* ZRK32 grew much better when cultivated in a rich medium compared with strains cultivated in the basal medium (Figure 2A). We also found that *N*-acetyl glucosamine did not stimulate the growth of strain ZRK32. These findings have revealed that strain ZRK32 prefers a nutrient-rich medium, which is different from most other *Planctomycetes* bacteria (Wiegand et al., 2018; Wiegand et al., 2020b).

Notably, when growing in the rich medium, the expressions of most genes involved in the TCA cycle and EMP glycolysis pathway in strain ZRK32 were upregulated (Figure 2B-D, Figure S5B, and Figure S6), suggesting that strain ZRK32 might function through the complete TCA metabolic pathway and EMP glycolysis pathway to obtain energy for growth (Figure S8) (Zheng et al., 2021b). Consistent with the presence of EMP glycolysis pathway in strain ZRK32, we found that it could use a variety of sugars including glucose, maltose, fructose, isomaltose, galactose, D-mannose, and rhamnose (Table S2). As for the presence of TCA cycle in the anaerobic strain ZRK32, we propose that it might use other alternative electron acceptors (such as sulfate reducers, nitrate reducers, iron reducers, etc) in place of oxygen for the TCA cycle, as shown in other anaerobic bacteria (Alteri et al., 2012).

Strain ZRK32 possesses a condensed and intact nucleoid-like structure and a complex membrane structure (Figure 1D), which is similar to other reported *Planctomycetes* bacteria (Kaushik et al., 2020; Dedysh et al., 2021). In addition, TEM observation of ultrathin sections of cells from strain ZRK32 showed some eukaryote-like structures (Figure S9, panels 1 to 6). Although it is impossible to judge accurately what these structures are based on current methods, the internal structures of *Planctomycetes* bacteria are nevertheless fascinating. We also observed some vesicle-like (Figure S9, panels 7 to 8) and vacuole-like structures (Figure S9, panel 9) in strain ZRK32, similar to those observed in other *Planctomycetes* members (Wiegand et al., 2018). Vacuoles store cellular components (such as proteins and sugars etc.,) and play essential roles in plant responses to different biotic/abiotic signaling pathways (Zhang et al., 2014). The presence of vacuoles in strain ZRK32 suggests that *Planctomycetes* bacteria might have adopted a eukaryotic mechanism for nutrient metabolism and signal transduction.

Most species within the class *Planctomycetia* divide by budding, and species within the class *Phycisphaerae* divide by binary fission (Fukunaga et al., 2009; Yoon et al., 2014; Kovaleva et al., 2015; Kallscheuer et al., 2020a; Pradel et al., 2020; Salbreiter et al., 2020; Wiegand et al., 2020b). However, the *Poriferisphaera* strain ZRK32 (Figure 1C and Figure S1) was demonstrated to divide by budding, similar to the *Poriferisphaera* strain KS4 proposed by Kallscheuer et *al* (2020) (Kallscheuer et al., 2020b), suggesting that members within the class *Phycisphaerae* might divide through both binary fission and budding. It is noteworthy that *P. heterotrophicis* ZRK32 forms an empty cell framework first, followed by the entry of the cellular contents (Figure 3A, B), which is different from the typical budding mode observed in yeast, where the cellular framework and contents both extend simultaneously (Herskowitz, 1988). During microbial cell division, members of the Fts family of proteins (tubulin homologs) usually assemble at the future site of cell division, forming a contractile ring known as the Z ring (Wiegand et al., 2018). We observed that the expressions of numerous Fts-related proteins (FtsJ, FtsK, FtsL, FtsW, FtsX, and FtsY) in strain ZRK32 were upregulated when the strain had been cultivated in the rich medium (Figure 3C). However, the FtsZ protein was absent from strain ZRK32, which is consistent with the proposal that the *ftsZ* gene is absent from *Planctomycetes* genomes (Jogler et al., 2012). In addition, we also found that some genes that encoded rod shape-determining proteins (MreB, MreC, and MreD) were present in strain ZRK32, and that their expressions were upregulated in the strains cultivated in the rich medium (Figure 3C). MreB, usually participates in the formation and degradation of peptidoglycan, which ultimately determines bacterial cell shape (Rohs and Bernhardt, 2021).

*Planctomycetes* bacteria are major players in the global nitrogen cycling and perform important reactions, such as the anaerobic ammonium oxidation process that oxidizes NH_4_^+^ to N_2_ gas, using NO_2_^-^ as an electron acceptor (Strous et al., 1998; Oshiki et al., 2016; Wiegand et al., 2018). To date, most studies on anaerobic ammonium oxidation have been related to the monophyletic group (*Candidatus* Brocadiae) in the phylum *Planctomycetes* (Strous et al., 1999), with only few reports on the process of nitrogen metabolism in other *Planctomycetes* bacteria. Here, we found that NO_3_^-^ and NH_4_^+^ promoted the growth of strain ZRK32, while NO_2_^-^ inhibited its growth (Figure 4A). Based on our data and results from others, we speculate that NO_3_^-^ might act as a terminal electron acceptor in the respiratory electron transport chain of strain ZRK32. In addition, NO_2_^-^ might react with the iron-sulfur proteins in strain ZRK32 to form iron-nitric oxide complexes, which then inactivate iron-sulfur enzymes and inhibit the growth of strain ZRK32 (Reddy et al., 1983). We speculate that strain ZRK32 converts NO_3_^-^ to NH_4_^+^ (Figure 4B, C), enabling the NH_4_^+^ created to enter the glutamate metabolic pathway (Figure 4D, E) — a pathway that is closely associated with several processes, including nitrogen metabolism, the TCA cycle, the EMP glycolysis pathway, and amino acid metabolism (Figure S8). Consistently, in the presence of NO_3_^-^ or NH_4_^+^, genes associated with the TCA cycle and the EMP glycolysis pathway were upregulated (Figure 4F, Figure S5C and Figure S7). The presence of rich nutrients, and either NO_3_^-^ or NH_4_^+^ also stimulated the expression of genes encoding the NADH-ubiquinone oxidoreductase complex (Figures 2D and 4G). This complex couples the oxidation of NADH and the reduction of ubiquinone to generate a proton gradient, which is then used for ATP synthesis (Reda et al., 2008). Notably, large number of genes associated with flagellum assembly were also up-regulated in strain ZRK32 (Figures 2E and 4H). Flagellum-mediated motility is beneficial for bacteria, not only because it allows them to respond quickly to an ever-changing environment but also because it enables them to seek and acquire nutrients for survival (Wadhams and Armitage, 2004; Zheng et al., 2021a). Thus, strain ZRK32 might regulate the formation and motility of its flagellum to accelerate the absorption and utilization of nutrients available in its environment.

One of our most exciting results was that NO_3_^-^ or NH_4_^+^ could induce the release of a chronic bacteriophage (Phage-ZRK32) from strain ZRK32 (Figure 5A). The Phage-ZRK32 was capable of facilitating nitrogen metabolism and amino acid metabolism in strain ZRK32 through the function of AMGs, which incorporate nitrogen into certain amino acids (including glutamate, cysteine and glycine) (Figure 5B) (Orlowski and Meister, 1970; Mouilleron and Golinelli-Pimpaneau, 2007; Iyer et al., 2009; Lu, 2013). Phylogenetic analyses were performed (Figures S10-13) using AMG amino acid sequences (including amidoligase, glutamine amidotransferase, gamma-glutamylcyclotransferase, and glutathione synthase) from Phage-ZRK32, in addition to the same amino acid sequences from related phages and their bacterial hosts. The results showed that these AMGs may have been acquired from *Pseudomonas* via horizontal gene transfer (Matilla et al., 2014; Zhou et al., 2019). Consistently, Phage-ZRK32 promoted the growth of a marine *Pseudomonas* bacterium (*Pseudomonas stutzeri* 273) in the presence of either NO_3_^-^ or NH_4_^+^ (Figure 5C, D). Given that the AMGs encoding amidoligase and gamma-glutamylcyclotransferase were absent in the genome of *Pseudomonas stutzeri* 273, we speculate that Phage-ZRK32 might promote the growth of *Pseudomonas stutzeri* 273 by facilitating nitrogen metabolism and amino acid generation, as in the *Planctomycetes* strain ZRK32. Most bacteriophage life-cycles are described as a lytic or lysogenic cycle (Du Toit, 2017). Currently, more attention has been given to the chronic life cycle, where bacterial growth continues despite phage reproduction (Hoffmann Berling and Maze, 1964), and the progeny of these phage particles are released from host cells via extrusion or budding without killing the host (Putzrath and Maniloff, 1977; Russel, 1991; Marvin et al., 2014). Undoubtedly, Phage-ZRK32, which was induced by either NO_3_^-^ or NH_4_^+^ in *Planctomycetes* strain ZRK32, is a chronic bacteriophage. Moreover, it has recently been reported that the tailless *Caudoviricetes* phage particles are enclosed in lipid membrane and are released from the host cells by a nonlytic mechanism (Liu et al., 2022), and the prophage induction contributes to the production of membrane vesicles by *Lacticaseibacillus casei* BL23 during cell growth (da Silva Barreira et al., 2022). Considering that strain ZRK32 has a large number of membrane vesicles during cell growth (Figure S9), we speculated that Phage-ZRK32 might be a membrane vesicle-engulfed phage and its release should be related to membrane vesicles. Altogether, our findings provide a novel insight into nitrogen metabolism in *Planctomycetes* bacteria and provide a suitable model to study the interactions between *Planctomycetes* and viruses.

## Materials and Methods

### Enrichment and cultivation of deep-sea *Planctomycetes* bacteria

To isolate and cultivate *Planctomycetes* bacteria, 2 g deep-sea sediment samples were added to a 500 mL anaerobic bottle containing 400 mL basal medium (1.0 g/L yeast extract, 20.0 g/L NaCl, 1.0 g/L CH_3_COONa, 1.0 g/L NaHCO_3_, 0.5 g/L KH_2_PO_4_, 0.2 g/L MgSO_4_.7H_2_O, 0.7 g/L cysteine hydrochloride, 500 µL/L 0.1 % (w/v) resazurin, 1.0 L sterilized distilled water, pH 7.0) supplemented with 1.0 g/L NH_4_Cl and 1.0 g/L NaNO_3_, ensuring headspace volume was retained above the liquid surface. The medium was prepared under a 100% N_2_ gas phase and sterilized by autoclaving at 115 °C for 30 minutes; following this, rifampicin (100 µg/mL) was added. The inoculated media were anaerobically incubated at either 4 °C or 28 °C for one month. The basal medium supplemented with 1.0 g/L NH_4_Cl, 1.0 g/L NaNO_3_, and 15 g/L agar was evenly spread on to the inside wall of a Hungate tube, which formed a thin layer of medium for the bacteria to grow. After this, 50 µL of the enriched culture was anaerobically transferred into an anaerobic roll tube and then spread on to the medium layer. These tubes were also anaerobically cultured at either 4 °C or 28 °C for ten days. Single colonies growing at 28 °C were selected using sterilized bamboo sticks; they were then cultured in the 15 mL Hungate tube containing 10 mL basal medium (supplemented with 1.0 g/L NH_4_Cl and 1.0 g/L NaNO_3_) at 28 °C for seven days under a 100% N_2_ atmosphere. One strain was identified as a member of the phylum *Planctomycetes*, but was noted to have less than 98% 16S rRNA gene sequence similarity to other cultured strains; this strain was therefore selected to be cultivated. Strain ZRK32 was selected and purified by repeating the Hungate roll-tube method. The purity of strain ZRK32 was confirmed regularly by observation using a transmission electron microscope (TEM) and by repeating partial sequencing of the 16S rRNA gene. As strain ZRK32 grew slowly in basal medium, we used a rich culture medium (10.0 g/L yeast extract, 20.0 g/L NaCl, 1.0 g/L CH_3_COONa, 1.0 g/L NaHCO_3_, 0.5 g/L KH_2_PO_4_, 0.2 g/L MgSO_4_.7H_2_O, 0.7 g/L cysteine hydrochloride, 500 µL/L 0.1 % (w/v) resazurin, 1.0 L sterilized distilled water, pH 7.0).

### TEM observation

To observe the morphological characteristics of strain ZRK32, 10 mL culture was collected by centrifuging at 5000 × *g* for ten minutes. Cells were then washed three times with PBS buffer (137 mM NaCl, 2.7 mM KCl, 10 mM Na_2_HPO_4_, 1.8 mM KH_2_PO_4_, 1 L sterile water, pH 7.4). Finally, the cells were suspended in 20 μL PBS buffer, and then transferred onto copper grids coated with a carbon film by immersing the grids in the cell suspension for 30 minutes (Zheng et al., 2021b). To observe the ultrastructure of strain ZRK32, ultrathin-sections were prepared using methods previously described (Graham and Orenstein, 2007). Briefly, 500 mL of cells (cultured for six days at 28 °C) were collected by centrifuging at 5000 × *g* for 20 minutes, and then washing three times with PBS buffer. The cells were then preserved in 2.5% (v/v) glutaraldehyde for 12 hours at 4 °C and then dehydrated using different ethanol concentrations (30%, 50%, 70%, 90%, and 100%) for ten minutes each time. The cells were then embedded in a plastic resin. Finally, 50-70 nm ultrathin sections were produced using an ultramicrotome (Leica EM UC7, Germany) and then stained using uranyl acetate and lead citrate. All samples were examined under TEM (HT7700, Hitachi, Japan).

### Phylogenetic analysis

To construct a maximum likelihood 16S rRNA phylogenetic tree, the full-length 16S rRNA gene sequences of strain ZRK32 and other related taxa were extracted from their corresponding genomes (www.ncbi.nlm.nih.gov/). The maximum likelihood genome phylogenetic tree was constructed from a concatenated alignment of 37 protein-coding genes (Wu et al., 2013) (extracted from the genomes using Phylosift v1.0.1) (Darling et al., 2014); all genes were present in a single copy and were universally distributed in both archaea and bacteria (Table S3). The reference sequences of four AMGs encoding amidoligase, glutamine amidotransferase, gamma-glutamylcyclotransferase, and glutathione synthase were retrieved by blasting the phage gene against the entire NCBI database, respectively. All phylogenetic trees were constructed using the W-IQ-TREE web server (http://iqtree.cibiv.univie.ac.at) (Trifinopoulos et al., 2016) using the “GTR+F+I+G4” model, and the Interactive Tree of Life (iTOL v5) online tool (Letunic and Bork, 2021) was used to edit the phylogenetic trees.

### Growth assays of strain ZRK32

To assess the effects of nutrient-rich media on strain ZRK32 growth, we set up different cultures using either rich medium or basal medium. To assess the effects of different inorganic nitrogen sources (20 mM NO_3_^-^, 20 mM NH_4_^+^, and 20 mM NO_2_^-^) on strain ZRK32 growth, we used a rich culture medium supplemented with the nitrogen sources mentioned above. For each growth assay, 15 mL of strain ZRK32 culture was inoculated in a 2LJL Hungate bottle containing 1.5LJL of the respective media. All Hungate bottles were anaerobically incubated at 28 °C. Bacterial growth was monitored by measuring daily OD_600_ values via a microplate reader until cell growth reached a stationary phase. Three replicates were performed for each condition. The concentrations of NO_3_ ^-^, NH_4_ ^+^, and NO_2_ ^-^ were determined using a continuous flow analyzer (SKALAR-SAN++, Netherlands), which has an analytical precision of 6%.

### Transcriptomics

For transcriptomic sequencing, strains of ZRK32 were cultured in 1.5 L of either basal medium, rich medium, or rich medium supplemented with different nitrogen sources (20 mM NO_3_^-^, 20 mM NH_4_^+^, or 20 mM NO_2_ ^-^) at 28 °C for six days. Three biological replicates were cultured for each condition. Following this, cells from the three replicates were mixed and then collected for transcriptomic sequencing by Novogene (Tianjin, China), as previously described (Zheng et al., 2021b; Zheng et al., 2022). Detailed protocols describing library preparation, clustering, sequencing, and data analyses are described in the supplementary information. All heat maps were made by HemI 1.0.3.3.

### Isolation of bacteriophages

Isolation of the bacteriophages was performed using similar methods to those described previously, but with some modifications (Yamamoto et al., 1970; Tseng et al., 1990; Kim and Blaschek, 1991). Strain ZRK32 was inoculated in either rich medium, or rich medium supplemented with 20 mM NO_3_^-^ or 20 mM NH_4_^+^, and then incubated at 28 °C for six days. Different cultures were collected by centrifuging at 8000 × *g* at 4 °C for 20 minutes; this was repeated three times. The supernatant was filtered through a 0.22 μm millipore filter (Pall Supor, USA), and then 1 M NaCl was added to lyse the residual bacteria. The supernatant was collected by centrifuging at 8000 × *g* at 4 °C for 20 minutes. The phage particles were immediately precipitated with 100 g/L polyethylene glycol (PEG8000) at 4 °C for two hours, and collected by centrifuging at 10000 × *g* at 4 °C for 20 minutes. The phage particles were then suspended in SM buffer (0.01% gelatin, 50 mM Tris-HCl, 100 mM NaCl, and 10 mM MgSO_4_). The suspension was then extracted three times using an equal volume of chloroform (Lin et al., 2012) and collected by centrifuging at 4000 × *g* at 4 °C for 20 minutes. Finally, the clean phage particles were obtained. Detailed protocols describing the genome sequencing of the bacteriophages are described in the supplementary information.

### Growth assay of *Pseudomonas stutzeri* 273 cultured in oligotrophic medium supplemented with different nitrogen sources and Phage-ZRK32

The assistance role of the bacteriophage induced from strain ZRK32 was tested in another deep-sea bacterium *Pseudomonas stutzeri* 273 (Wu et al., 2016). Specifically, 50 µL freshly incubated *Pseudomonas stutzeri* 273 cells were inoculated in 5 mL of either oligotrophic medium (10 g/L NaCl, 0.1 g/L yeast extract, 1 L sterilized distilled water), oligotrophic medium supplemented with 20 µl/L Phage-ZRK32 (without the extraction by chloroform), oligotrophic medium supplemented with 20 mM NO ^-^ or NH_4_^+^, oligotrophic medium supplemented with 20 mM of either NO_3_^-^ or NH_4_^+^, or 20 µl/L Phage-ZRK32. The cultures were then incubated for 24 hours at 28 °C. Three biological replicates for each culture condition were performed. The progress of the bacterial growth was monitored by measuring OD_600_ values using a microplate reader (Infinite M1000 Pro, Switzerland) every four hours until cell growth reached a stationary phase.

### Data availability

The full-length 16S rRNA gene sequence of strain ZRK32 has been deposited at GenBank under the accession number MW376756. The complete genome sequence of strain ZRK32 has been deposited at GenBank under the accession number CP066225. The raw sequencing reads from the transcriptomics analyses of ZRK32 strains cultured with different concentrations of yeast extract and nitrogen sources have been deposited into the NCBI Short Read Archive (accession numbers: PRJNA768630 and PRJNA694614, respectively). The genome sequence of Phage-ZRK32 has been deposited into the GenBank database with the accession number OP650935.

## Acknowledgements

This work was funded by the Major Research Plan of the National Natural Science Foundation (Grant No. 92051107), the NSFC Innovative Group Grant (No. 42221005), Science and Technology Innovation Project of Laoshan Laboratory (Grant No. LSKJ202203103; 2022QNLM030004-3), Shandong Provincial Natural Science Foundation (ZR2021ZD28), Strategic Priority Research Program of the Chinese Academy of Sciences (Grant No. XDA22050301), China Ocean Mineral Resources R&D Association Grant (Grant No. DY135-B2-14), Key Collaborative Research Program of the Alliance of International Science Organizations (Grant No. ANSO-CR-KP-2022-08), Key deployment projects of Center of Ocean Mega-Science of the Chinese Academy of Sciences (Grant No. COMS2020Q04), and the Taishan Scholars Program (Grant No. tstp20230637) for Chaomin Sun. We thank Dr. Diana Walsh from Life Science Editors for her great effort to improve the writing quality of our manuscript.

## Author contributions

RZ, CW, and CS conceived and designed the study; RZ and CW conducted most of the experiments; RL collected the samples from the deep-sea cold seep; RC helped to construct evolutionary trees; RZ, CW, and CS lead the writing of the manuscript; all authors contributed to and reviewed the manuscript.

