## Supplementary methods, figures and tables for "Physiological and metabolic insights into the first cultured anaerobic representative of deep-sea *Planctomycetes* bacteria"

### Supplementary Results

#### Genomic and physiologic analyses of strain ZRK32

To understand more characteristics of strain ZRK32, its whole genome was sequenced and analyzed. The genome size of strain ZRK32 was 5,234,020 bp with a DNA G+C content of 46.28 mol% (Figure. S3). Annotation of the genome of strain ZRK32 revealed that it consisted of 4,175 predicted genes including 6 rRNA genes (2, 2, and 2 for 5S, 16S, and 23S, respectively) and 45 tRNA genes, which were higher than those reported in the most closely related type strain *Poriferisphaera corsica* KS4$^T$ (Table S1). Moreover, the genome size (5,234,020 bp) and gene numbers (4,175) of strain ZRK32 were also higher than those in strain KS4$^T$ (4,291,168 bp, 3,714). Strain ZRK32 was able to grow over a temperature range of 4-32 ℃ (optimum, 28 ℃), which was wider than that of strain KS4$^T$ (15-30 ℃, optimum 27 ℃) (Figure. S4A). The pH range for growth of strain ZRK32 was 6.0-8.0 (optimum, pH 7.0) (Figure. S4B). Growth of strain ZRK32 was observed at 0.5-5.0% NaCl (Figure. S4C).

#### Description of *Poriferisphaera heterotrophicis* sp. nov.

*Poriferisphaera heterotrophicis* (hetero'tro.phicis. L. fem. adj. *heterotrophicis* means a heterotrophic lifestyle). Cells are spherical, average diameter of 0.4-1.0 µm, strictly anaerobic and have a single polar flagellum. The temperature range for growth is 4-32 ℃ with an optimum at 28 ℃. Growing at pH values of 6.0-8.0 (optimum, pH 7.0). Growth occurs at NaCl concentrations from 0.5% to 5.0%. The type strain, ZRK32$^T$, was isolated from a deep-sea cold seep sediment, P. R. China. The DNA G+C content of the type strain is 46.28 mol%.

### Supplementary Methods

#### Physiological tests

Effects of temperature, pH, and NaCl concentration on the growth of strain ZRK32 were determined in the rich medium as described above. To evaluate the temperature range for growth, cultures were incubated at 4, 16, 24, 28, 32, 37, 45, 60 °C (pH 7.0).

To determine the pH range for growth, the medium was adjusted at optimum temperature (28 ℃) to pH 4.0-10.0 with increments of 0.5 pH units under a 100% $N_2$ atmosphere. NaCl requirements were tested in the modified rich medium (without 20.0 g/L NaCl) supplemented with 0-10% (w/v) NaCl (1.0% intervals). Single sugar (including glucose, maltose, fructose, sucrose, starch, isomaltose, trehalose, galactose, cellulose, xylose, D-mannose, and rhamnose) was added from sterile filtered stock solutions to the final concentration at 20 mM, respectively. Cell culture containing only 0.02 g yeast extract ($L^{-1}$) without adding any other substrates was used as a control. These cultures were incubated at 28 °C for 14 days and then the $OD_{600}$ values were measured via a microplate reader (Infinite M1000 Pro; Tecan, Mannedorf, Switzerland). For each experiment, three biological replicates were performed.

**Genome sequencing, annotation, and analysis of strain ZRK32**

For genomic sequencing, strain ZRK32 was grown in the liquid rich medium and harvested after one week of incubation at 28 ℃. Genomic DNA was isolated by using the PowerSoil DNA isolation kit (Mo Bio Laboratories Inc., Carlsbad, CA). Thereafter, the genome sequencing was carried out with both the Illumina NovaSeq PE150 (San Diego, USA) and Nanopore PromethION platform (Oxford, UK) at the Beijing Novogene Bioinformatics Technology Co., Ltd. A complete description of the library construction, sequencing, and assembly was performed as previously described (Zheng et al., 2021). We used seven databases to predict gene functions, including Pfam (Protein Families Database, http://pfam.xfam.org/), GO (Gene Ontology, http://geneontology.org/) (Ashburner et al., 2000), KEGG (Kyoto Encyclopedia of Genes and Genomes, http://www.genome.jp/kegg/) (Kanehisa et al., 2004), COG (Clusters of Orthologous Groups, http://www.ncbi.nlm.nih.gov/COG/) (Galperin et al., 2015), NR (Non-Redundant Protein Database databases), TCDB (Transporter Classification Database), and Swiss-Prot (http://www.ebi.ac.uk/uniprot/) (Bairoch and Apweiler, 2000). A whole genome Blast search (E-value less than 1e-5, minimal alignment length percentage larger than 40%) was performed against above seven databases.

In addition, the genome relatedness values were calculated by multiple approaches, including Average Nucleotide Identity (ANI) based on the MUMMER ultra-rapid aligning tool (ANIm) and the BLASTN algorithm (ANIb), the tetranucleotide signatures (Tetra), and *in silico* DNA-DNA (*is*DDH) similarity. ANIm, ANIb, and Tetra values were calculated using the JSpecies WS (http://jspecies.ribohost.com/jspeciesws/) (Richter et al., 2016). The recommended species criterion cut-offs were used: 95% for the ANIb and ANIm, 0.99 for the Tetra signature. The *is*DDH similarity values were calculated by the Genome-to-Genome Distance Calculator (GGDC) (http://ggdc.dsmz.de/) (Meier-Kolthoff et al., 2013). A value of 70% *is*DDH similarity was used as a recommended standard for delineating species.

**The detailed procedure for transcriptomic sequencing analysis of strain ZRK32 cultured under different conditions.**

**(1) Library preparation for strand-specific transcriptome sequencing.** A total amount of 3 μg RNA per sample was used as input material for the RNA sample preparation. Sequencing libraries were generated using NEBNext® Ultra™ Directional RNA Library Prep Kit for Illumina® (NEB, USA) following the manufacturer's recommendations and index codes were added to attribute sequences to each sample. Then, rRNA was removed using a specialized kit that left the mRNA. Fragmentation was carried out using divalent cations under elevated temperature in NEBNext First Strand Synthesis Reaction Buffer (5×). First strand cDNA was synthesized using random hexamer primer and M-MuLV Reverse Transcriptase (RNaseH$^-$). Second strand cDNA synthesis was subsequently performed using DNA Polymerase I and RNase H. In the reaction buffer, dNTPs with dTTP were replaced by dUTP. Remaining overhangs were converted into blunt ends via exonuclease/polymerase activities. After adenylation of 3' ends of DNA fragments, NEBNext Adaptor with hairpin loop structure was ligated to prepare for hybridization.

In order to select cDNA fragments of preferentially 150~200 bp in length, the library fragments were purified with AMPure XP system (Beckman Coulter, USA). Then 3 μL USER Enzyme (NEB，USA) was used with size-selected, adaptor-ligated cDNA at 37 ℃ for 15 min followed by 5 min at 95 ℃ before PCR. PCR was performed with Phusion High-Fidelity DNA polymerase, Universal PCR primers, and Index (X) Primer. At last, products were purified (AMPure XP system) and library quality was assessed on the Agilent Bioanalyzer 2100 system.

**(2) Clustering and sequencing.** The clustering of the index-coded samples was performed on a cBot Cluster Generation System using TruSeq PE Cluster Kit v3-cBot-HS (Illumia) according to the manufacturer's instructions. After cluster generation, the library preparations were sequenced on an Illumina Hiseq platform and paired-end reads were generated.

**(3) Data analysis.** Raw data of fastq format were firstly processed through in-house perl scripts. In this step, clean data were obtained by removing reads containing adapter, reads containing ploy-N and low quality reads from raw data. At the same time, Q20, Q30, and GC content the clean data were calculated. All the downstream analyses were based on the clean data with high quality. Reference genome and gene model annotation files were downloaded from genome website directly. Both building index of reference genome and aligning clean reads to reference genome were used Bowtie2-2.2.3 (setting: -D 15 -R 2 -N 0 -L 22 -i S,1,1.15) (Langmead and Salzberg, 2012). HTSeq v0.6.1 (default parameters) was used to count the reads numbers mapped to each gene. FPKM of each gene was calculated based on the length of the gene and reads count mapped to this gene. FPKM, expected number of Fragments Per Kilobase of transcript sequence per Millions base pairs sequenced, considers the effect of sequencing depth and gene length for the reads count at the same time, and is currently the most commonly used method for estimating gene expression levels (Trapnell et al., 2009).

**(4) Differential expression analysis.** Differential expression analysis was performed using the DESeq R package (1.18.0) and edgeR v3.24.3 (|$\log_2$(Fold change)| >= 1 & padj <= 0.05) (Anders and Huber, 2010). DESeq provide statistical routines for determining differential expression in digital gene expression data using a model based on the negative binomial distribution. The resulting *P*-values were adjusted using the Benjamini and Hochberg's approach for controlling the false discovery rate. Genes with an adjusted *P*-value < 0.05 found by DESeq were assigned as differentially expressed. (For DEGSeq without biological replicates) Prior to differential gene expression analysis, for each sequenced library, the read counts were adjusted by edgeR program package through one scaling normalized factor. Differential expression analysis of two conditions was performed using the DEGSeq R package (1.20.0) (Wang et al., 2010). The *P* values were adjusted using the Benjamini & Hochberg method. Corrected *P*-value of 0.005 and $\log_2$ (Fold change) of 1 were set as the threshold for significantly differential expression.

**(5) GO and KEGG enrichment analysis of differentially expressed genes.** Gene Ontology (GO) enrichment analysis of differentially expressed genes was implemented by the GOseq R package, in which gene length bias was corrected (Young et al., 2010). GO terms with corrected *P* value less than 0.05 were considered significantly enriched by differential expressed genes. KEGG is a database resource for understanding high-level functions and utilities of the biological system, such as the cell, the organism, and the ecosystem, from molecular-level information, especially large-scale molecular datasets generated by genome sequencing and other high-throughput experimental technologies (http://www.genome.jp/kegg/) (Kanehisa et al., 2008). We used KOBAS software to test the statistical enrichment of differential expression genes in KEGG pathways.

**Real-Time Quantitative Reverse Transcription PCR (qRT-PCR).**

To validate the RNA-seq data, we determined the expression levels of some genes by qRT-PCR. For qRT-PCR, cells of strain ZRK32 cultured in 1.5 L of either basal medium, rich medium, or rich medium supplemented with different nitrogen sources (20 mM $NO_3^-$, 20 mM $NH_4^+$ or 20 mM $NO_2^-$, respectively) at 28 °C for six days were collected at 8000 × *g* for 20 minutes. Three biological replicates were cultured for each condition. Total RNA from each sample was extracted using the Trizol reagent (Solarbio, China). The RNA concentration was measured using Spectrophotometer (NanoPhotometer NP80, Implen, Germany). Then RNAs from corresponding samples were reverse transcribed into cDNA (complementary DNA) using ReverTra Ace™ qPCR RT Master Mix with gDNA Remover (TOYOBO, Japan). The transcriptional levels of different genes were determined by qRT-PCR using SYBR® Green Realtime PCR Master Mix (TOYOBO, Japan) and the QuantStudio™ 6 Flex (Thermo Fisher Scientific, USA). The PCR condition was set as following: initial denaturation at 95 ℃ for 3 min, followed by 40 cycles of denaturation at 95 ℃ for 10 s, annealing at 56 ℃ for 20 s, and extension at 72 ℃ for 20 s. The 16S rRNA gene of strain ZRK32 was used as an internal reference and the gene expression was calculated using the $2^{-\Delta\Delta Ct}$ method (Livak and Schmittgen, 2001), with each transcript signal normalized to that of 16S rRNA gene. Transcript signals for each treatment were compared to those of control group. Specific primers for genes associated with the TCA cycle, NADH-ubiquinone oxidoreductase, flagellum assembly, and EMP glycolysis of strain ZRK32 and 16S rRNA gene were designed using Primer 5.0 as shown in Table S4. All qRT-PCR runs were conducted with three biological and three technical replicates.

**A detailed procedure for genome sequencing analysis of phages**

To sequence the genome of bacteriophage, the phage genomic DNA was extracted from different purified phage particles. Firstly, to remove residual host DNA, 1 μg/mL DNase I and RNase A were added to the concentrated phage solution for nucleic acid digestion overnight at 37 ℃. The digestion treatment was inactivated at 80 ℃ for 15 min, followed by extraction with a Viral DNA Kit (Omega Bio-tek, USA) according to the manufacturer's instructions. Then, the genome sequencing was performed by

Biozeron Biological Technology Co.Ltd (Shanghai, China). The detailed process of library construction, sequencing, genome assembly, and annotation was described below.

**(1) Library construction and Illumina HiSeq sequencing.** Briefly, for Illumina pair-end sequencing of each phage, 0.2 μg genomic DNA was used for the sequencing library construction. Paired-end libraries with insert sizes of ~400 bp were prepared following the standard procedure. The purified genomic DNA was sheared into smaller fragments with a desired size by Covaris, and blunt ends were generated using the T4 DNA polymerase. The desired fragments were purified through gel-electrophoresis, then enriched and amplified by PCR. The index tag was introduced into the adapter at the PCR stage and we performed a library quality test. Finally, the qualified Illumina pair-end library was used for Illumina NovaSeq 6000 sequencing (150 bp*2, Shanghai BIOZERON Co., Ltd).

**(2) Genome assembly.** The raw paired end reads were trimmed and quality controlled by the Trimmomatic (version 0.36, http://www.usadellab.org/cms/uploads/supplementary/Trimmomatic) (Pollet et al., 2011) with parameters (SLIDINGWINDOW: 4:15, MINLEN: 75). Clean data were obtained and used for further analysis. We have used the ABySS software (http://www.bcgsc.ca/platform/bioinfo/software/abyss) to perform genome assembly with multiple-Kmer parameters and got the optimal results. The GapCloser software (https://sourceforge.net/projects/soapdenovo2/files/GapCloser/) was subsequently applied to fill up the remaining local inner gaps and correct the single base polymorphism for the final assembly results.

**(3) Genome Annotation.** For bacteriophages, these obtained genome sequences were subsequently annotated by searching these predicted genes against non-redundant (NR in NCBI, 20180814), SwissProt (release-2021_03, http://uniprot.org) (Dedysh and Ivanova, 2019), KEGG (Release 94.0, http://www.genome.jp/kegg/) (Buckley et al., 2006), COG (update-2020_03, http://www.ncbi.nlm.nih.gov/COG) (Brümmer et al., 2004), and CAZy (update-2021_09, http://www.cazy.org/) (Woebken et al., 2007) databases.

### Supplementary Figures

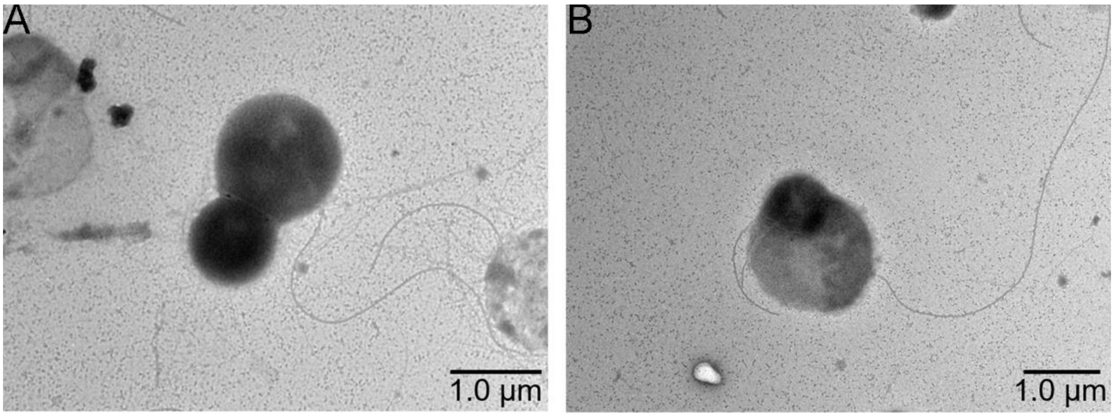


**Figure S1. TEM observation the morphology of cells from *P. heterotrophicis* ZRK32.**

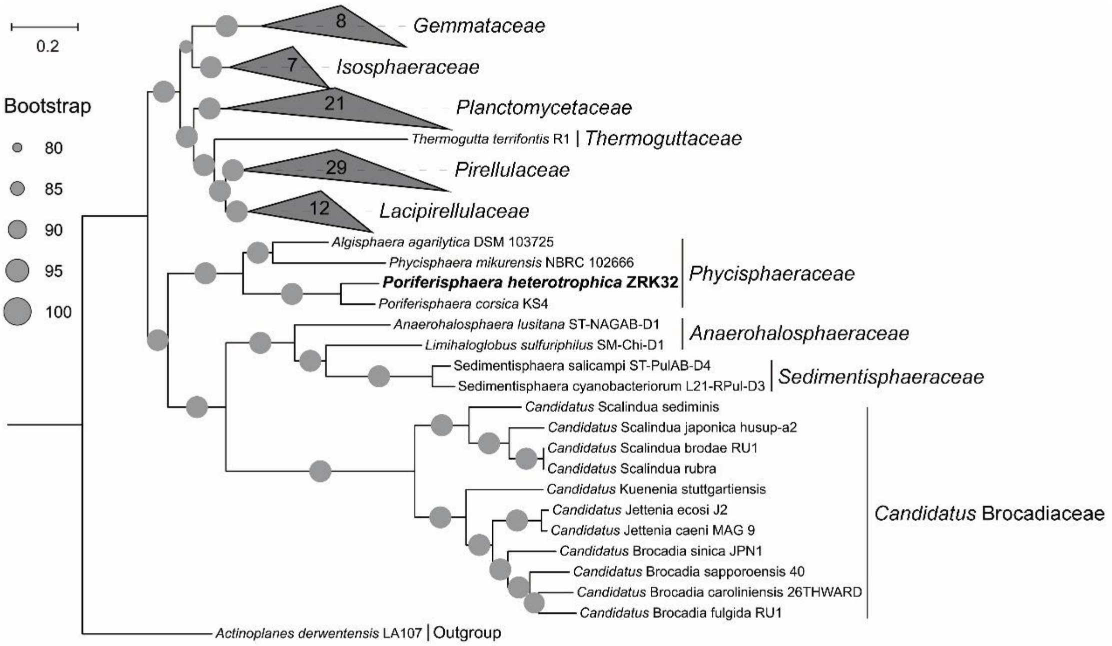


**Figure S2. Maximum likelihood phylogenetic tree of genome sequences from the *P. heterotrophicis* ZRK32 and other *Planctomycetes* bacteria constructed from the concatenated alignment of 37 single-copy genes; *Actinoplanes derwentensis* LA107 was used as the outgroup.** The tree was inferred and reconstructed using the maximum likelihood criterion, with bootstrap values (%) > 80; these are indicated at the base of each node with a gray dot (expressed as a percentage from 1,000 replications). Bar, 0.2 substitutions per nucleotide position.

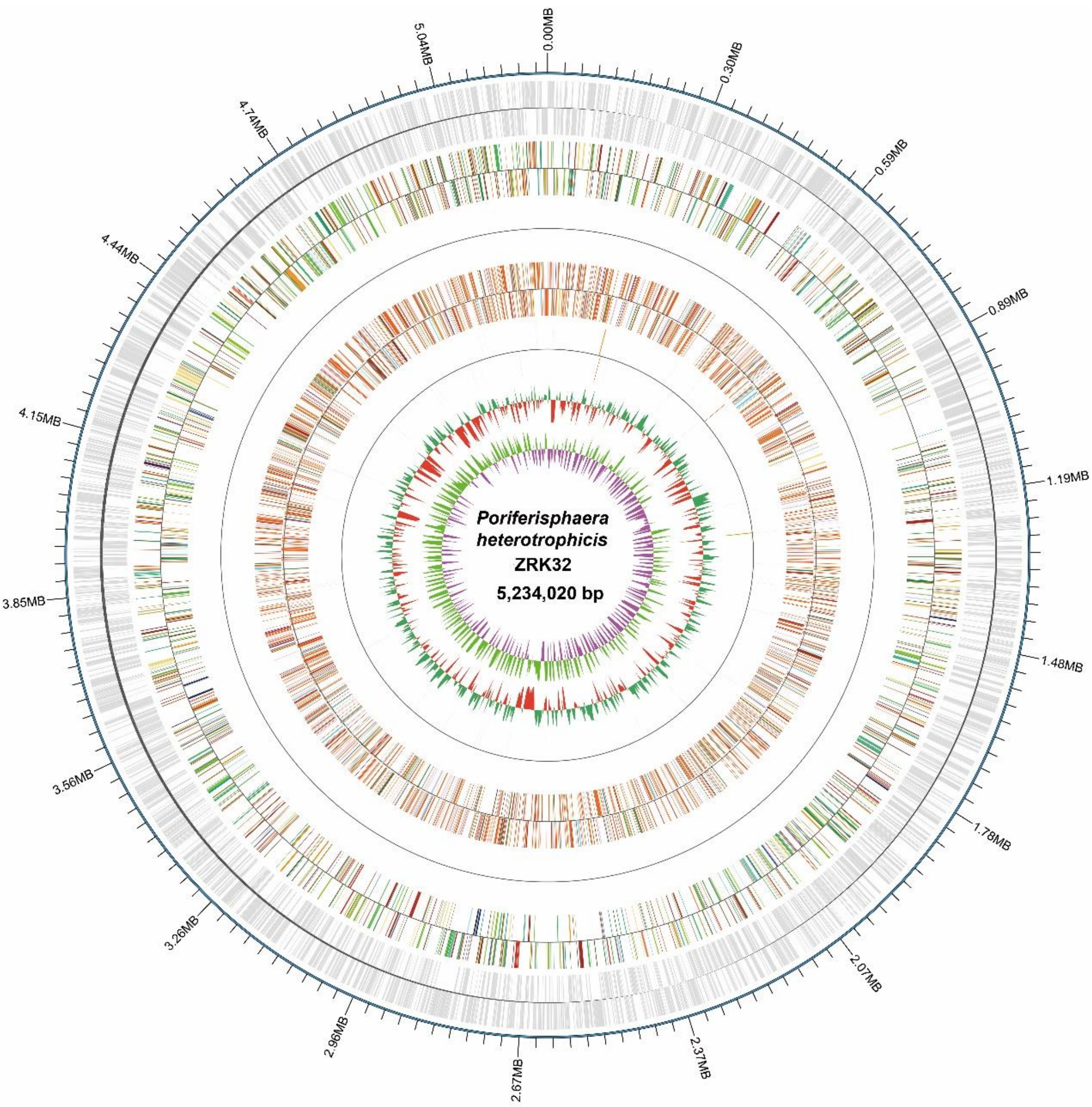


**Figure S3. Circular diagram of the *P. heterotrophicis* ZRK32 genome.** Rings indicate, from outside to the center: a genome-wide marker with a scale of 0.3 MB; coding genes; gene function annotation results; ncRNA; GC content; GC skew.

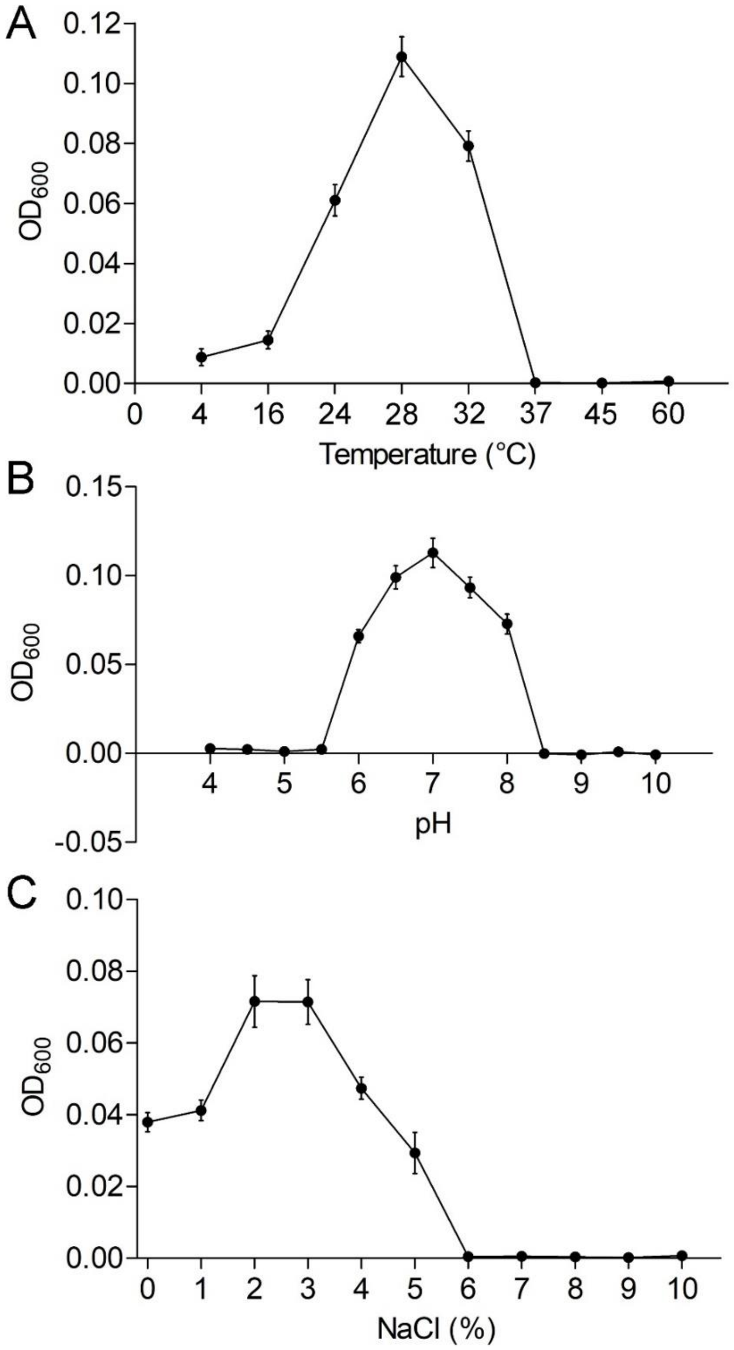


**Figure S4. Physiological characterizations of *P. heterotrophicis* ZRK32.** Growth curves of ZRK32 strains cultivated in different conditions. Temperature (A), pH (B), and NaCl concentration (C) ranges enabling growth were analyzed of ZRK32 strains cultivated in rich medium with three biological triplicates.

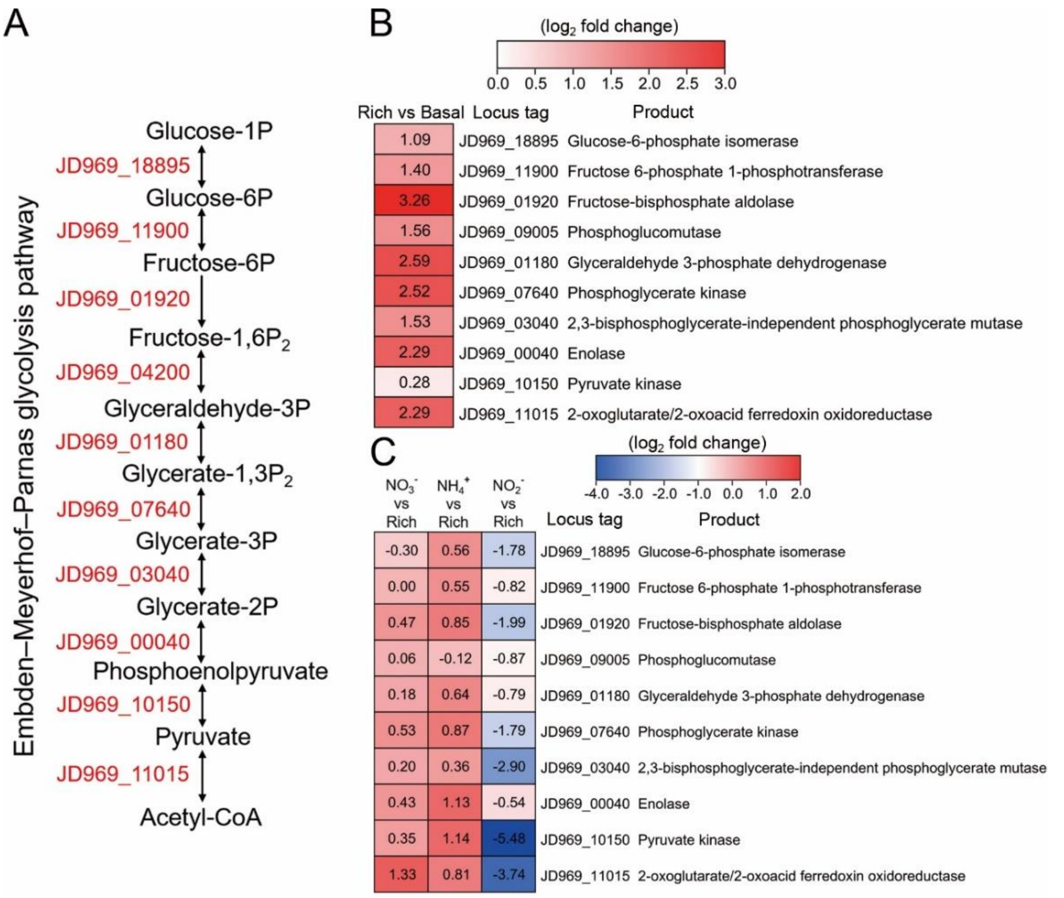


**Figure S5. Transcriptomics analysis of the genes associated with the EMP glycolysis pathway of *P. heterotrophicis* ZRK32 strains cultivated in the rich medium alone and cultivated in rich medium supplemented with either 20 mM $NO_3^-$, 20 mM $NH_4^+$, or 20 mM $NO_2^-$.** (A) Diagram of the EMP glycolysis pathway. The gene numbers shown in this schemeatic are the same as those shown in panels B and C. (B) Transcriptomics-based heat map showing differentially expressed genes associated with the EMP glycolysis pathway of strain ZRK32 cultivated in rich medium (Rich) compared with strains cutivated in basal medium (Basal). (C) Transcriptomics-based heat map showing the relative expression levels of genes associated with the EMP glycolysis pathway of strain ZRK32 cultivated in the rich medium supplemented with different inorganic nitrogen sources (20 mM $NO_3^-$, 20 mM $NH_4^+$ or 20 mM $NO_2^-$) compared with strains cultivated in the rich medium alone. "Rich" indicates rich medium. "$NO_3^-$, $NH_4^+$, and $NO_2^-$" indicate rich medium supplemented with 20 mM $NO_3^-$, 20 mM $NH_4^+$, and 20 mM $NO_2^-$, respectively. The numbers in panels B and C represent the fold change of gene expression (by using the $log_2$ value).

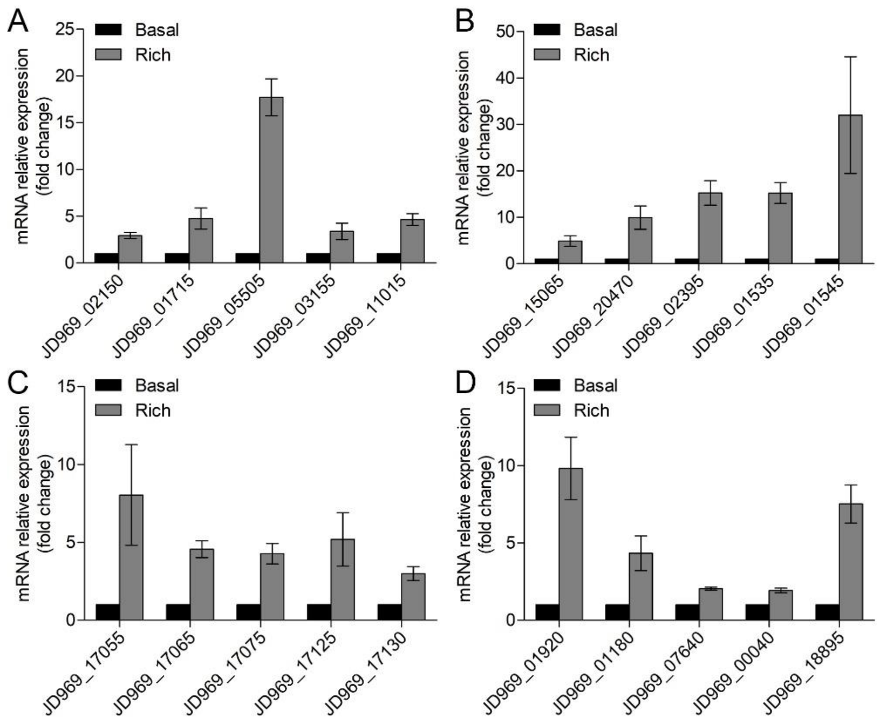


**Figure S6. qRT-PCR detection of the relative expression levels of the genes associated with the TCA cycle (A), NADH-quinone oxidoreductase (B), flagellar assembly (C), and EMP glycolysis pathway (D) of *P. heterotrophicis* ZRK32 strains cultivated in rich medium (Rich) compared with strains cultivated in basal medium (Basal).** JD969_02150, succinate dehydrogenase; JD969_01715, fumarate hydratase; JD969_05505, malate dehydrogenase; JD969_03155, isocitrate dehydrogenase; JD969_11015, 2-oxoglutarate ferredoxin oxidoreductase; JD969_15065, NADH-quinone oxidoreductase subunit A; JD969_20470, NADH-quinone oxidoreductase subunit C; JD969_02395, NADH-quinone oxidoreductase subunit F; JD969_01535, NADH-quinone oxidoreductase subunit L; JD969_01545, NADH-quinone oxidoreductase subunit N; JD969_17055, flagellar basal body rod protein FlgB; JD969_17065, flagellar hook-basal body protein FliE; JD969_17075, flagellar motor switch protein FliG; JD969_17125, motility protein MotA; JD969_17130, motility protein MotB; JD969_01920, fructose-bisphosphate aldolase; JD969_01180, glyceraldehyde 3-phosphate dehydrogenase; JD969_07640, phosphoglycerate kinase; JD969_00040, enolase; JD969_18895, glucose-6-phosphate isomerase.

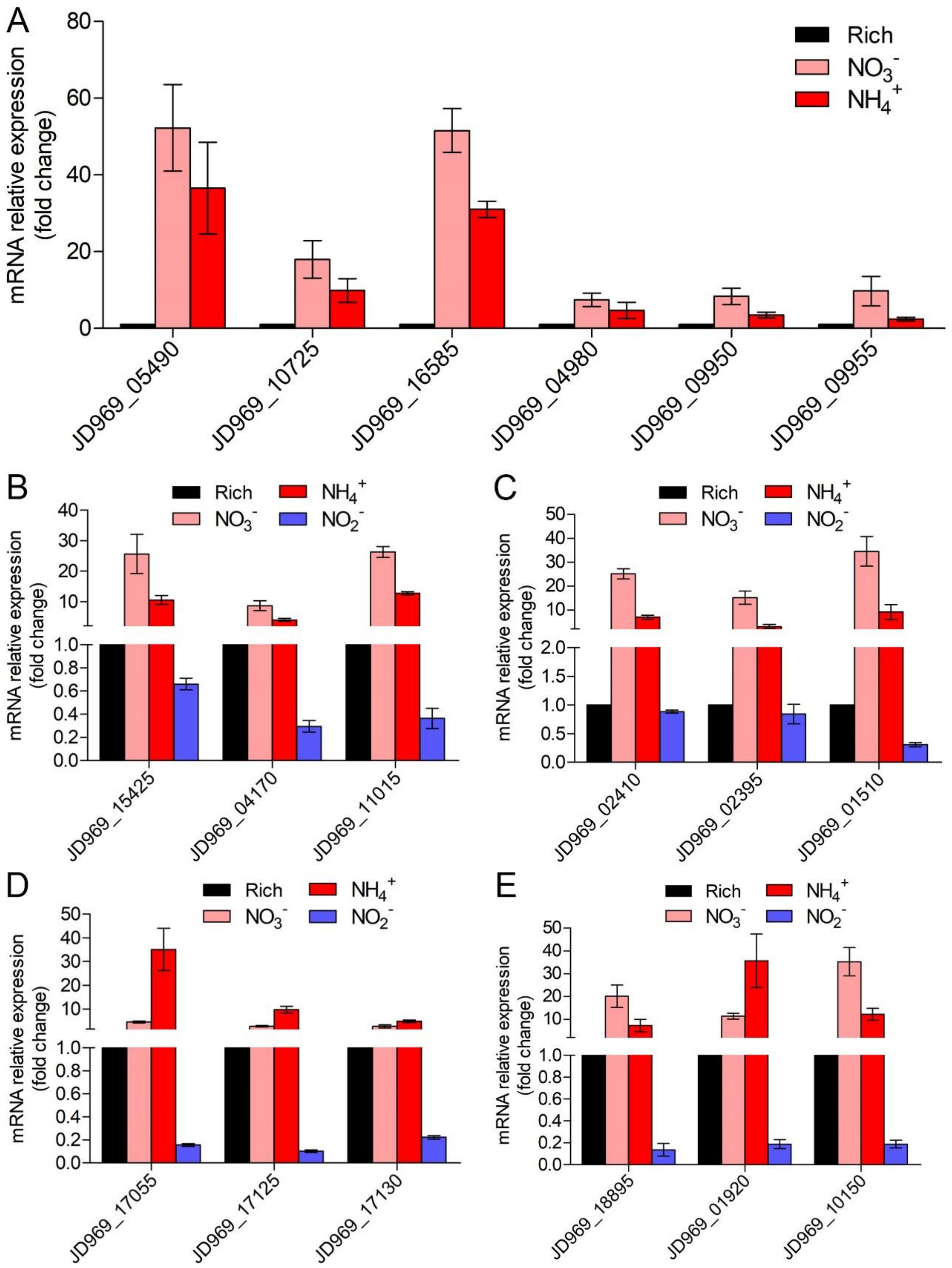


**Figure S7. qRT-PCR detection of the relative expression levels of the genes associated with nitrogen metabolism (A), TCA cycle (B), NADH-quinone oxidoreductase (C), flagellar assembly (D), and EMP glycolysis pathway (E) of ZRK32 strains cultivated in the rich medium supplemented with different inorganic nitrogen sources (20 mM $NO_3^-$, 20 mM $NH_4^+$ or 20 mM $NO_2^-$)**

**compared with strains cultivated in the rich medium alone.** "Rich" indicates rich medium. "$NO_3^-$, $NH_4^+$, and $NO_2^-$" indicate rich medium supplemented with 20 mM $NO_3^-$, 20 mM $NH_4^+$, and 20 mM $NO_2^-$, respectively. JD969_05490, nitrate reductase; JD969_10725, nitrate reductase; JD969_16585, nitrite reductase; JD969_04980, glutamine synthetase; JD969_09950, glutamate synthase; JD969_09955, glutamate synthase; JD969_15425, citrate synthase; JD969_04170, aconitate hydratase; JD969_11015, 2-oxoglutarate ferredoxin oxidoreductase; JD969_02410, NADH-quinone oxidoreductase subunit D; JD969_02395, NADH-quinone oxidoreductase subunit F; JD969_01510, NADH-quinone oxidoreductase subunit G; JD969_17055, flagellar basal body rod protein FlgB; JD969_17125, motility protein MotA; JD969_17130, motility protein MotB; JD969_18895, glucose-6-phosphate isomerase; JD969_01920, fructose-bisphosphate aldolase; JD969_10150, pyruvate kinase.

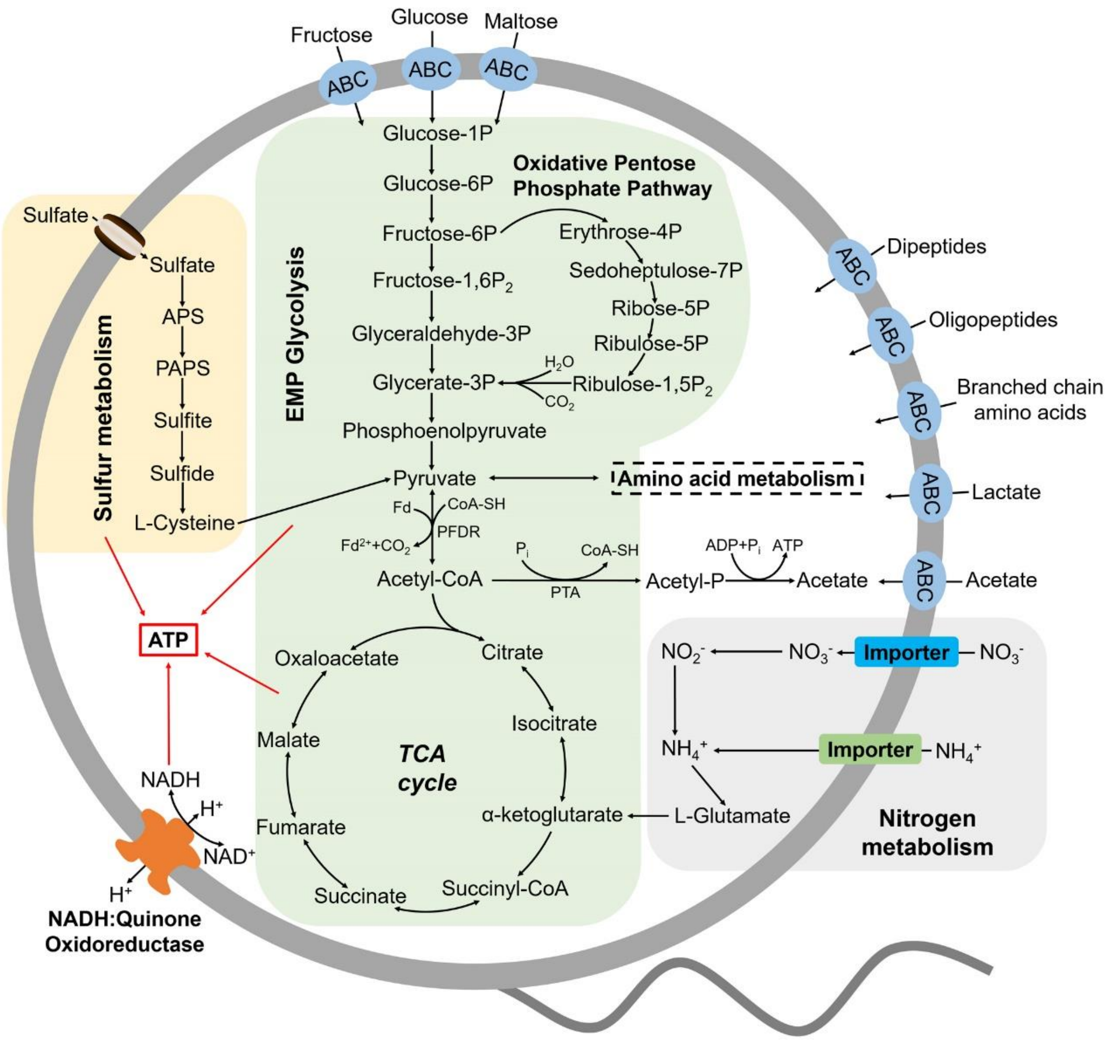


**Figure S8. Muti-omics based central metabolism model of *P. heterotrophicis* ZRK32.** Based on the combination of genomic, transcriptomic and physiological characteristics, we proposed a model towards the central metabolic traits of strain ZRK32. In this model, central metabolisms including the EMP glycolysis pathway, the oxidative pentose phosphate pathway, the TCA cycle, sulfur metabolism, nitrogen metabolism and electron transport system were shown. All the above items are closely related to the energy production in strain ZRK32. Briefly, strain ZRK32 contains a number of genes related to ABC transporters of amino acids and peptides, which could transport these organic matters into the cell to participate in the EMP glycolysis and oxidative pentose phosphate pathway. These processes eventually drive the formation of pyruvate and acetyl-CoA, which enter the TCA cycle to produce energy for the growth of strain ZRK32. Moreover, nitrate could be converted to ammonium through the dissimilatory nitrate reduction, which participates in the synthesis of L-Glutamate and thereby entering into the TCA cycle for energy generation. Meanwhile, the $H^+$-transporting NADH: Quinone oxidoreductase required for energy production is present in strain ZRK32. Strain ZRK32 also contains a complete pathway for assimilatory sulfate reduction, which contributes to to energy production.

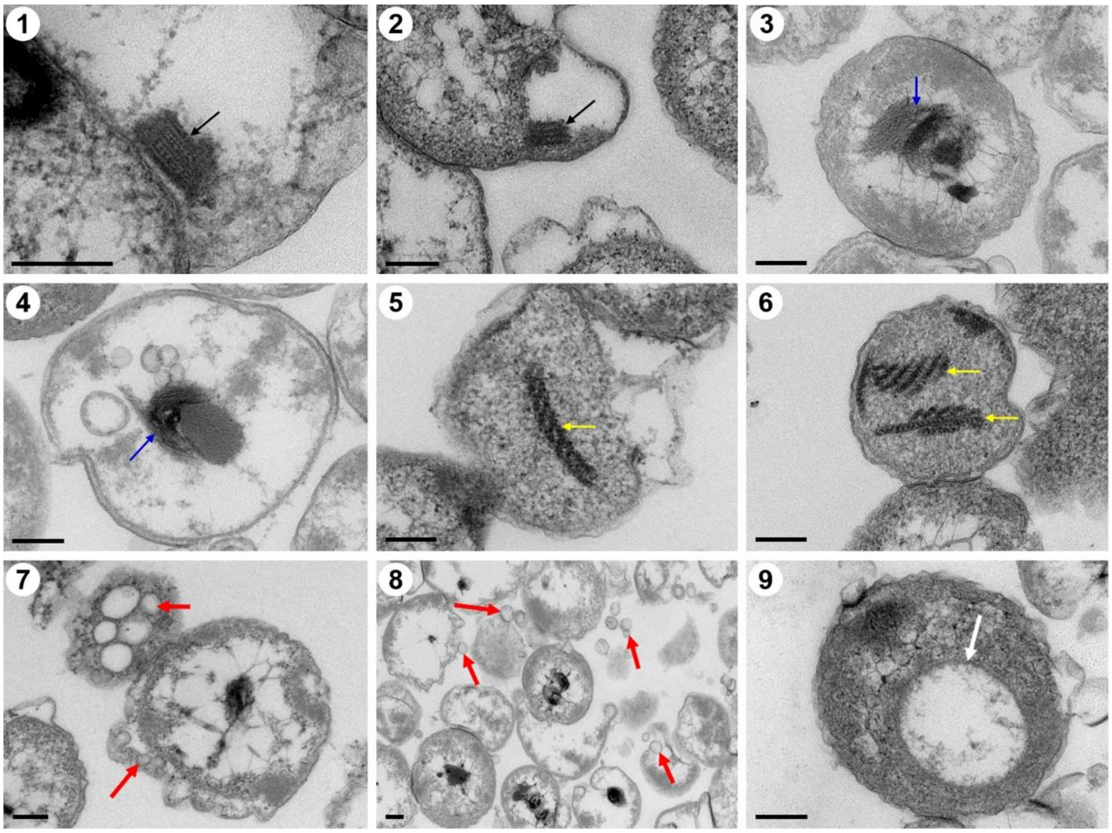


**Figure S9. Ultrathin TEM sections showing some eukaryote-like structures observed in cells from *P. heterotrophicis* ZRK32.** Bars: 200 nm.

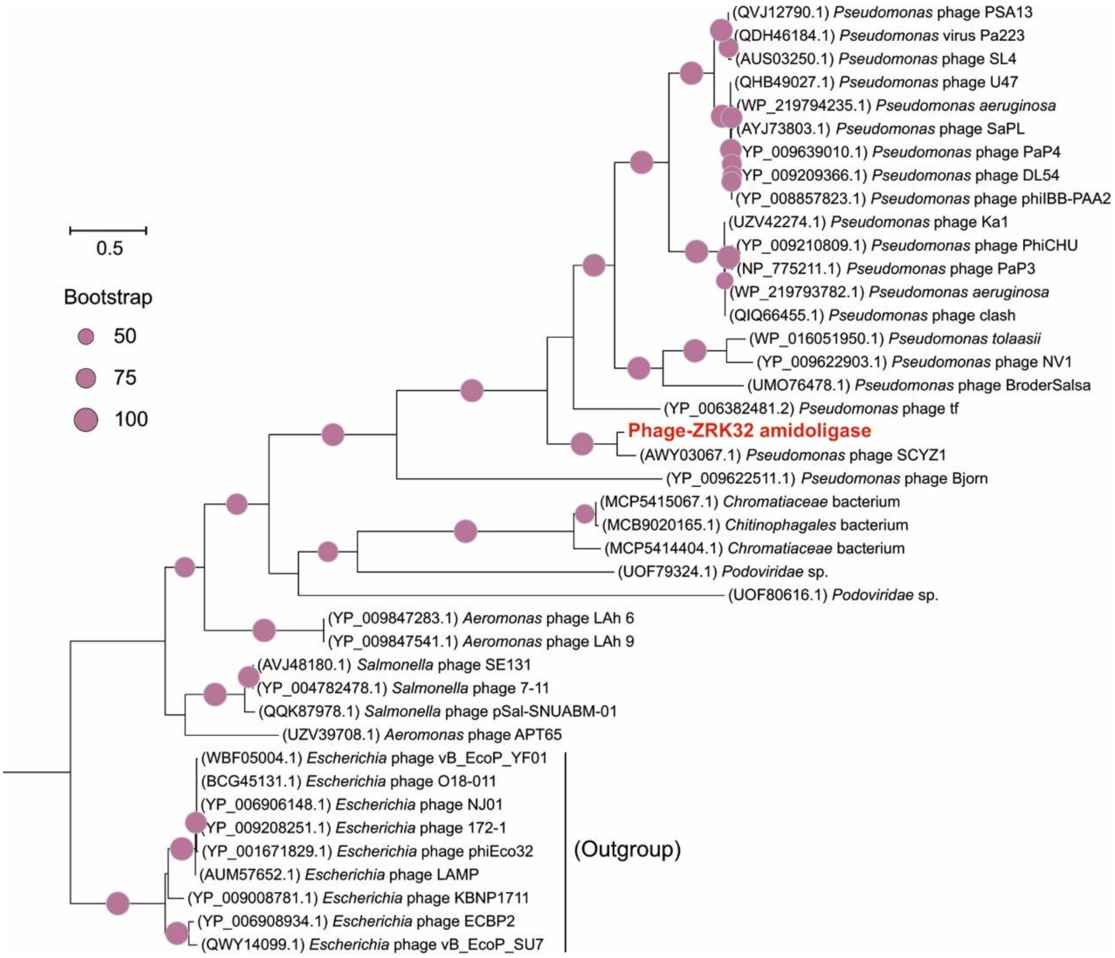


**Figure S10. Phylogenetic analysis of Phage-ZRK32, some related phages, and bacterial hosts, based on the aligned amino acid sequences of amidoligase.** The NCBI accession number for each amino acid sequence is indicated after each corresponding strain's name. The amino acid sequences of amidoligase from nine *Escherichia* phages were used as the outgroup. The tree was inferred and reconstructed using the maximum likelihood criterion, with bootstrap values (%) > 50; these are indicated at the base of each node with a gray dot (expressed as a percentage from 1,000 replications). Bar, 0.5 substitutions per nucleotide position.

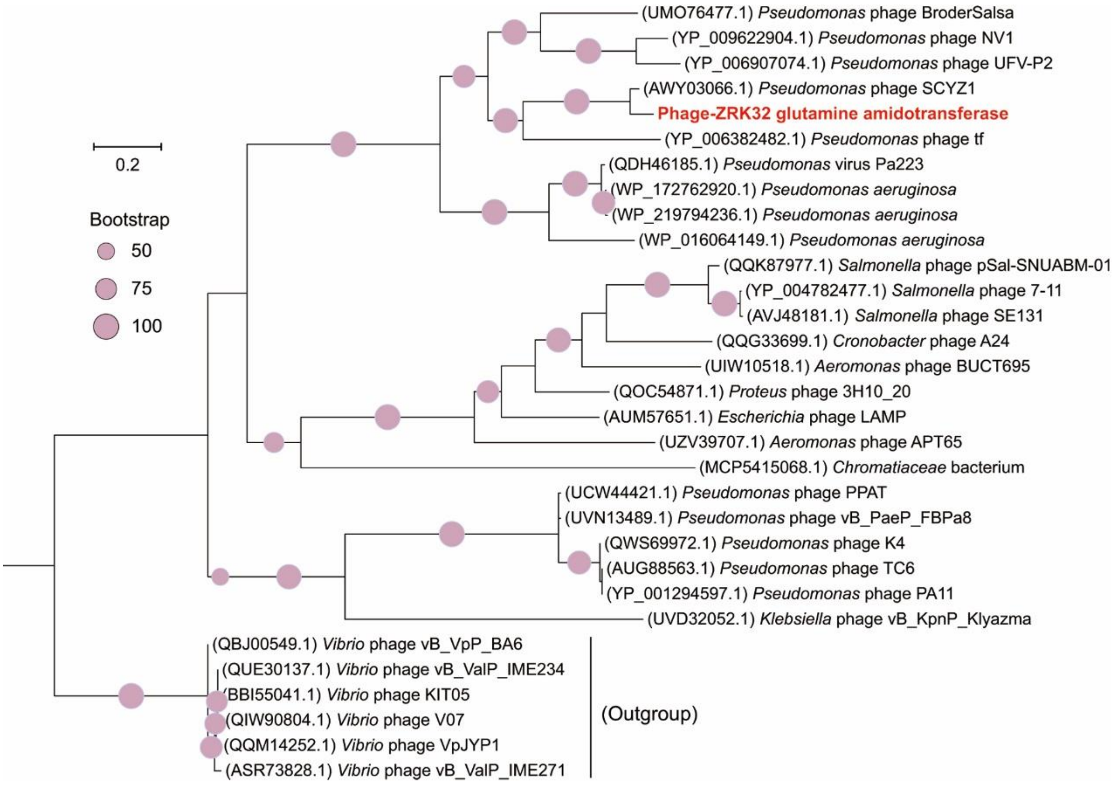


**Figure S11. Phylogenetic analysis of Phage-ZRK32, some related phages, and bacterial hosts, based on the aligned amino acid sequences of glutamine amidotransferase.** The NCBI accession number for each amino acid sequence is indicated after each corresponding strain's name. The amino acid sequences of glutamine amidotransferase from six *Vibrio* phages were used as the outgroup. The tree was inferred and reconstructed using the maximum likelihood criterion, with bootstrap values (%) > 50; these are indicated at the base of each node with a gray dot (expressed as a percentage from 1,000 replications). Bar, 0.2 substitutions per nucleotide position.

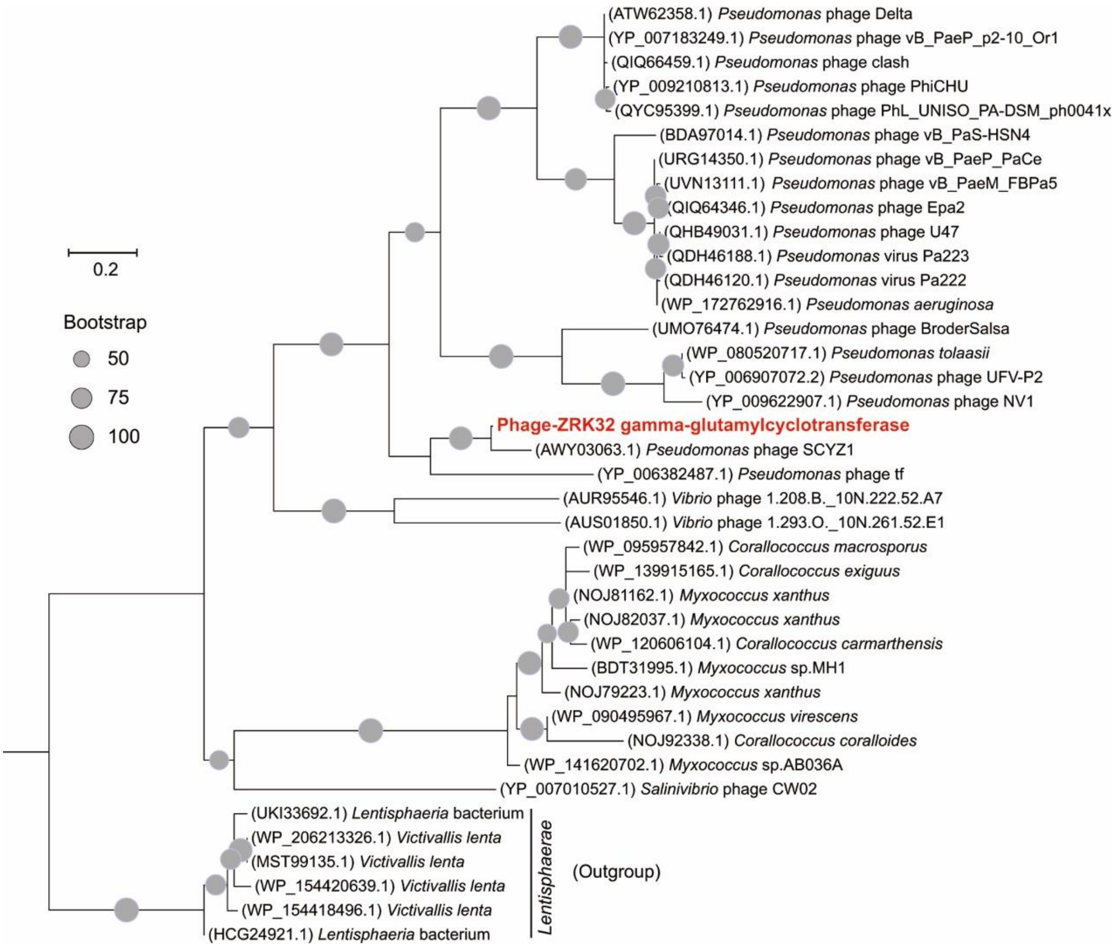


**Figure S12. Phylogenetic analysis of Phage-ZRK32, some related phages, and bacterial hosts, based on the aligned amino acid sequences of gamma-glutamylcyclotransferase.** The NCBI accession number for each amino acid sequence is indicated after each corresponding strain's name. The amino acid sequences of gamma-glutamylcyclotransferase from six *Lentisphaerae* strains were used as the outgroup. The tree was inferred and reconstructed using the maximum likelihood criterion, with bootstrap values (%) > 50; these are indicated at the base of each node with a gray dot (expressed as a percentage from 1,000 replications). Bar, 0.2 substitutions per nucleotide position.

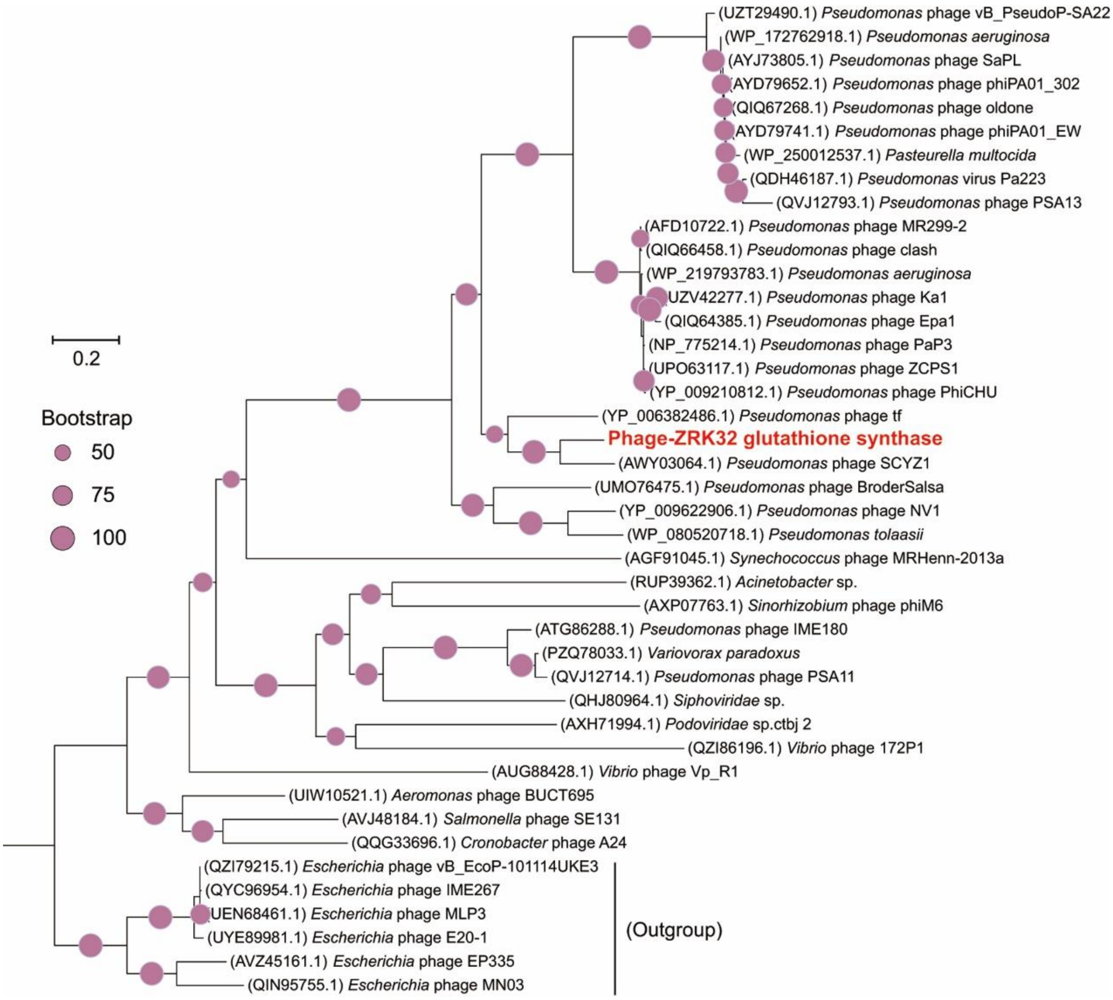


**Figure S13. Phylogenetic analysis of Phage-ZRK32, some related phages, and bacterial hosts, based on the aligned amino acid sequences of glutathione synthase.** The NCBI accession number for each amino acid sequence is indicated after each corresponding strain's name. The amino acid sequences of glutathione synthase from six *Escherichia* phages were used as the outgroup. The tree was inferred and reconstructed using the maximum likelihood criterion, with bootstrap values (%) > 50; these are indicated at the base of each node with a gray dot (expressed as a percentage from 1,000 replications). Bar, 0.2 substitutions per nucleotide position.

#### Supplementary Tables

**Table S1. Phenotypic and genotypic features of strain ZRK32 and the most closely related type strain *Poriferisphaera corsica* KS4$^{T}$.**

| Feature | ZRK32 | KS4$^{T}$ |
|---|---|---|
| **Phenotypic features** | | |
| Cell morphology | spherical | spherical |
| Cell diameter (μm) | 0.4-1.0 | 0.63 ±0.21 |
| Temperature range for growth (℃) | 4-32 | 15-30 |
| Optimum | 28 | 27 |
| pH for growth | 6.0-8.0 | 6.5-8.0 |
| Optimum | 7.0 | 7.5 |
| NaCl concentration range for growth (%) | 0-5 | 0-4 |
| Isolation source | Deep-sea cold seep sediment | Coast of the island Corsica |
| **Genomic features** | | |
| Gene Bank ID | CP066225 | CP036425 |
| Genome size (bp) | 5,234,020 | 4,291,168 |
| G+C content (mol%) | 46.28 | 48.7 |
| No.scaffolds/contigs | 1 | 1 |
| No. of genes | 4,175 | 3,714 |
| No. of rRNAs | 6 | 3 |
| No. of tRNAs | 45 | 45 |
| No. of protein-coding genes | 4,121 | 3,659 |
| Completeness (%) | 100 | 94.83 |
| ANIb (%) | 100 | 72.89 |
| ANIm (%) | 100 | 85.34 |
| Tetra | 1 | 0.97385 |
| *is*DDH (%) | 100 | 20.90 |

**Table S2. The sugar utilization of strain ZRK32.** +, Positive result or growth; −, negative result or no growth.

| **Sugar species** | **Utilization** |
|---|---|
| Glucose | + |
| Maltose | + |
| Fructose | + |
| Sucrose | - |
| Starch | - |
| Isomaltose | + |
| Trehalose | - |
| Galactose | + |
| Cellulose | - |
| Xylose | - |
| D-mannose | + |
| Rhamnose | + |

**Table S3. Marker genes used in the phylogenetic analysis.**

| ID | Protein |
|---|---|
| DNGNGWU00001 | ribosomal protein S2 rpsB |
| DNGNGWU00002 | ribosomal protein S10 rpsJ |
| DNGNGWU00003 | ribosomal protein L1 rplA |
| DNGNGWU00005 | translation initiation factor IF-2 |
| DNGNGWU00006 | metalloendopeptidase |
| DNGNGWU00007 | ribosomal protein L22 |
| DNGNGWU00009 | ribosomal protein L4/L1e rplD |
| DNGNGWU00010 | ribosomal protein L2 rplB |
| DNGNGWU00011 | ribosomal protein S9 rpsI |
| DNGNGWU00012 | ribosomal protein L3 rplC |
| DNGNGWU00013 | phenylalanyl-tRNA synthetase beta subunit |
| DNGNGWU00014 | ribosomal protein L14b/L23e rplN |
| DNGNGWU00015 | ribosomal protein S5 |
| DNGNGWU00016 | ribosomal protein S19 rpsS |
| DNGNGWU00017 | ribosomal protein S7 |
| DNGNGWU00018 | ribosomal protein L16/L10E rplP |
| DNGNGWU00019 | ribosomal protein S13 rpsM |
| DNGNGWU00020 | phenylalanyl-tRNA synthetase alpha subunit |
| DNGNGWU00021 | ribosomal protein L15 |
| DNGNGWU00022 | ribosomal protein L25/L23 |
| DNGNGWU00023 | ribosomal protein L6 rplF |
| DNGNGWU00024 | ribosomal protein L11 rplK |
| DNGNGWU00025 | ribosomal protein L5 rplE |
| DNGNGWU00026 | ribosomal protein S12/S23 |
| DNGNGWU00027 | ribosomal protein L29 |
| DNGNGWU00028 | ribosomal protein S3 rpsC |
| DNGNGWU00029 | ribosomal protein S11 rpsK |
| DNGNGWU00030 | ribosomal protein L10 |
| DNGNGWU00031 | ribosomal protein S8 |
| DNGNGWU00032 | tRNA pseudouridine synthase B |
| DNGNGWU00033 | ribosomal protein L18P/L5E |
| DNGNGWU00034 | ribosomal protein S15P/S13e |
| DNGNGWU00035 | Porphobilinogen deaminase |
| DNGNGWU00036 | ribosomal protein S17 |
| DNGNGWU00037 | ribosomal protein L13 rplM |
| DNGNGWU00039 | ribonuclease HII |
| DNGNGWU00040 | ribosomal protein L24 |

The DNGNGWU marker genes in phylosift refer to a suite of single-copy, protein-coding marker genes. All 37 DNGNGWU marker genes were concatenated to construct maximum likelihood phylogenetic tree.

**Table S4. Primers used for qRT-PCR.**

| Primer name | Nucleotide Sequence (5'-3') |
|---|---|
| 16S-F | CCCTTCCTTTGGGTCTGGTC |
| 16S-R | TTCCGCAATGCACGAAAGTG |
| JD969_02150-F | TACTGCTGAAGGCTTCCGTG |
| JD969_02150-R | ACGAGTTTGCTCGTCCAGAG |
| JD969_01715-F | CCAACTGCTGTACCACCGAT |
| JD969_01715-R | TGTTTGCTGGTTATGCACGC |
| JD969_05505-F | ATCAGCGAAGAACGCCTCAA |
| JD969_05505-R | TTCAGCCATCTGGACGGAAC |
| JD969_03155-F | TCCGCTACTTTGATGGCGTT |
| JD969_03155-R | TCGATGCCTGCGTAGATGTC |
| JD969_11015-F | GTTTGGCGAAGGCGTTTGAT |
| JD969_11015-R | TCCAGACCAACACTCAAGCC |
| JD969_15065-F | AGCGGAAGACACCTTTACGG |
| JD969_15065-R | GCACACAACTTGCCGATGTT |
| JD969_20470-F | GGGTGTCCATCGAAACGGAT |
| JD969_20470-R | CATCGAAGAAGACCCCGCAT |
| JD969_02395-F | AAGGAAGCCGGTACATGTGG |
| JD969_02395-R | GCCCAACCATCGTCAACAAC |
| JD969_01535-F | CTCACCGCTGGTTCAGTCAT |
| JD969_01535-R | AGCCGAAGAACATGAGCCAA |
| JD969_01545-F | TGCTCTGGGTTATCGCTGTC |
| JD969_01545-R | GTGAGCGACGGAGGAGTATG |
| JD969_17055-F | CGTCATCGTGTACTGGCAGA |
| JD969_17055-R | ACGTCGCTCACCAATAGCTC |
| JD969_17065-F | TTCAGCCTAAACAGGCGGTC |
| JD969_17065-R | GCATGCGATTCACTTGCTCA |
| JD969_17075-F | CCGCAGGCTTATGTTCGTCT |
| JD969_17075-R | GAGCCAAAGCCAATTCGTCG |
| JD969_17125-F | CACTCGCTGATCGTCTCGAA |
| JD969_17125-R | AACGCGAGGGTTATCACCAG |
| JD969_17130-F | CAGTTACGAACGTGCCAACG |
| JD969_17130-R | TCTGCAGTGTAAACGCGACT |

| | |
|---|---|
| JD969_01920-F | GGTAGACGCGTGGATCGTAG |
| JD969_01920-R | CACCGACACTCAGTACGCAT |
| JD969_01180-F | ATCGCTGGCATACTTGCTCA |
| JD969_01180-R | GGGTCACGTTCAGCACAAAC |
| JD969_07640-F | GCCGTCGGAGATATCGTTGT |
| JD969_07640-R | GGGTGGCGACAAAATCGTTC |
| JD969_00040-F | ACAGCGATACCGCTAACTCG |
| JD969_00040-R | CCTGAACGGTGTGAGAGGAC |
| JD969_18895-F | CCCACATTGGGAAACGCTTG |
| JD969_18895-R | ATTCGAACGTGCAGGCCTTA |
| JD969_10150-F | CAGCACACTCAACAACCTGC |
| JD969_10150-R | CGATGGCTTGGGGTTTTTCG |
| JD969_02410-F | ATACGCAATGTCGTGTGGGT |
| JD969_02410-R | TGCGAAACGACGCCTTATCT |
| JD969_01510-F | CAGCTCGCTACGATCGACAT |
| JD969_01510-R | AAGAGTGCGAGTACGAAGGC |
| JD969_15425-F | TCTTCGAGGCACCAGTCAAC |
| JD969_15425-R | CCGTGTTACGATATCCGGGG |
| JD969_04170-F | TGCTCACGCAACATCTCAGT |
| JD969_04170-R | TGATGCTTGGTCAGCCGATT |
| JD969_05490-F | GGTTTGCTGAATCATCCGCC |
| JD969_05490-R | CATTGCACGAAGTCAGCACC |
| JD969_10725-F | GTGTGGTATCCCGGCTTTGA |
| JD969_10725-R | TTCCTGGTGGCCGAATTGTT |
| JD969_16585-F | GAGTTAGATCCGGGTGCGAG |
| JD969_16585-R | AATCAGGCGGCCATTCTCAA |
| JD969_04980-F | CTCCAGGACGCACTCGATAC |
| JD969_04980-R | GACTGCGTTCATGAGTGGGA |
| JD969_09950-F | CTTGCACCGGGTATCACCTT |
| JD969_09950-R | CTTGCACCGGGTATCACCTT |
| JD969_09955-F | GTACCGGAAGAGCGTGTGAA |
| JD969_09955-R | TCAGACTGACAGAACGGCAC |
